# Gait as a Quantitative Translational Outcome Measure in Angelman Syndrome

**DOI:** 10.1101/2021.08.13.456146

**Authors:** Stela P. Petkova, Jessica D. Duis, Jill L. Silverman

## Abstract

Angelman Syndrome (AS) is a genetic neurodevelopmental disorder characterized by developmental delay, lack of speech, seizures, intellectual disability, and walking and balance disorders. Recently, motor ability became an interesting outcome measure in AS, as it is broad including ataxia, hypotonia, delayed and abnormal walking and postural movements and affects nearly every individual with AS. We predict that gait presents a strong opportunity for rigorous, reliable, and quantitative metrics with direct translation to evaluate pharmacological, dietary, and genetic therapies. Numerous motoric deficits have been identified clinically. In this study, we used an innovative, automated gait analysis as well as gold standard motor behavioral assays to further delineate components of motor, coordination, balance, and gait impairments in an AS mouse model across development. Our study demonstrated marked global motoric deficits in AS mice, corroborating many previous reports. Uniquely, this is the first report of nuanced and pertinent aberrations in quantitative spatial and temporal components of gait between AS and wildtype littermate controls, that are analogous in AS individuals. These metrics were followed longitudinally to observe the progression of maladaptive gait in AS, a clinical phenotype. This has not been reported previously and contributes a substantial novel metric for therapeutic development. Taken together, these findings demonstrate the robust translational value in the study of nuanced motor outcomes, i.e., gait, for AS, as well as similar genetic syndromes, in the endeavor of therapeutic screening.

**Lay Abstract:** Motor behaviors, like ambulation, gross and fine motor skills, coordination and balance, are easily quantifiable and readily translational between humans and preclinical rodent models for neurodevelopmental and neurodegenerative disorders, than other domains of behavior. To that end, we investigated gait across development in a mouse model for Angelman Syndrome and elucidated onset, progression, and decline of motor deficits in innovative, nuanced, and clinically relevant manner.

## Introduction

Angelman Syndrome (AS) is a rare genetic neurodevelopmental disorder (NDD) characterized by developmental delay; impaired receptive and expressive communication skills; motor disability; severe intellectual disabilities; and seizures (C. Williams & Franco, 2010; C. A. Williams, 2010; C. A. Williams, Driscoll, & Dagli, 2010). Movement disorders affect every individual with AS and are more prevalent than other commonly associated symptom. These movement abnormalities are not due to weakness or abnormal muscle tone, but may be accompanied by weakness or abnormal tone (Bird, 2014; Gentile et al., 2010; Schlaggar & Mink, 2003; Tan et al., 2011; Wheeler, Sacco, & Cabo, 2017). AS is caused by the loss of maternal expression of the gene *UBE3A* (ubiquitin-protein ligase E3A/E6AP) in the brain (Jiang et al., 1998; Matsuura et al., 1997; Sutcliffe et al., 1997). Due to brain-specific imprinting, the paternal allele is silenced; thus, loss of the maternal expression causes *UBE3A* deficiency isolated to the central nervous system (Sutcliffe et al., 1997; Yamasaki et al., 2003).

The AS motor profile includes gross and fine motor skill impairments; motor problems include spasticity, ataxia of gait, tremor, and muscle weakness. Delays in meeting motor milestones such neck control, limb coordination, and crawling are often noted in the first year of life, and additional motor deficits such as hypotonia, delays and impairments in independent walking abilities, and uncoordinated and ‘jerky’ movements are observed as children develop (Gentile et al., 2010; Micheletti et al., 2016; Wheeler et al., 2017). The delay in or lack of motor skill acquisition likely contributes to poor daily living skills, delay or failure to ambulate unassisted, and cognitive development observed later in life (Peters et al., 2004; Wheeler et al., 2017). Data from other neurodevelopmental disorders also show a strong relationship between the degree of motor deficit and other behavioral domains such as social communication and cognition (Champion et al., 2014; Gernsbacher et al., 2008; Leonard, 2016). Prior research in AS observed motor aberrations (DeLorey et al., 1998; Heck, Zhao, Roy, LeDoux, & Reiter, 2008; Jiang et al., 1998; Landers et al., 2005; Sinkkonen, Homanics, & Korpi, 2003; Sonzogni, Zhai, Mientjes, van Woerden, & Elgersma, 2020), without a direct examination of gait. Gait abnormalities such as ataxic gait with broad, instable stance have been well reported in AS children who walk independently, affecting upwards of 80% of patients in one report (Tan et al., 2011; Wheeler et al., 2017). To date, there is only one study that quantified gait, clinically, in AS (Grieco et al., 2018). Grieco et al. used the Zeno walkway, an electronic pressure mat covered with sensors which analyzes gait parameters, and found that AS children walked with small, fast steps. Moreover, at 6-9 years old, AS children had poor gait skills, more similar to typically developing children aged 1-3 years (Grieco et al., 2018).

*In vivo* models are essential for testing therapeutic delivery methods and preclinically and clinically relevant outcome measures are required to demonstrate the utility of innovative designs and validation of other traditional medicinal therapies that may be in the drug discovery pipeline for NDDs (Thurm, Kelleher, & Wheeler, 2020). Motor development is highly conserved across species, which is exemplified by gene expression profiles in the cortex and cerebellum (Strand et al., 2007). For humans and rodent models alike, motor deficits are the most consistently reported phenotype in AS. In mice, this behavioral phenotype has been well reported as reduced locomotor activity, poor balance and coordination, and impairment of performance on the accelerated rotarod (Born et al., 2017; Huang et al., 2013; Jiang et al., 1998; Miura et al., 2002). Recently, a novel rat model of rat with a full deletion of *Ube3a* recapitulated these motor deficits including the aforementioned accelerated rotarod motor learning deficit as well as additional nuanced impairments in rearing and fine forelimb motor skills (Berg et al., 2020; Dodge et al., 2020).

While there is a plethora of data on motor ability impairment, few studies tailor focus on gait and posture in behavioral reports in AS. The original report and behavioral phenotyping of an AS mouse used hind-paw footprint analysis to assess ataxia and noted a slightly reduced stride length in maternally-deficient *Ube3a* animals; footprint analysis was the original subjective, laborious, difficult to reproduce, garnering few indices of data method of gait collection. A later report, using the Dunnet et al. rudimentary paw painting down a runway of paper with manual scoring of metrics on both limbs, found elongated stride length and a wider stance only in the hindlimbs (Jiang et al., 1997; Heck et al., 2008). Herein, we applied treadmill walking, automated gait, longitudinally to further this line of inquiry, with enhanced rigor and reproducibility and observe and track development of motoric abnormalities in AS mice.

In this study, we focused on numerous of indices of gait, from physical metrics to temporal properties, analogs to clinical gait studies in AS, and nuanced approach to characterize gait patterns and discovered a maladaptive transition in gait in model mice. We compared the gait of AS mice and littermate controls across the lifespan to observe development, onset, progression, and severity of gait patterns. Our data revealed numerous significant alterations to gait in AS mice that are penetrant, detectable early in life, and robust, highlighting a potential quantitative biomarker that may be directly correlated with gait analysis in human subjects.

## Methods

### Animal Subjects

All animals were housed at the University of California Davis School of Medicine in Sacramento in a temperature-controlled vivarium on a 12:12 light-dark cycle. All procedures were conducted in compliance with the NIH Guidelines for the Care and Use of Laboratory Animals and approved by the Institutional Animal Care and Use Committee of UC Davis. A colony was maintained by breeding male B6.129S7-Ube3a^tm1Alb^/J (Jackson Laboratory, Stock No. 016590) with female C57B6/J mice, maintaining paternal transmission of the mutant allele. Angelman syndrome model mice used in behavioral experiments have maternal transmission of the mutant allele and were generated by crossing heterozygous dams with C57B6/J males to produce maternally inherited mutant *Ube3a*^m-/p+^ (AS, N=20) mice and wildtype littermate controls (WT, N=30). Mice were tested in the gait task from P22 (weaning) to P180 and motor assays were conducted at 4 months of age. Both sexes were used.

### Motor Behavior Assays

Motor behavior was tested at four months of age. Animals were habituated in a 30 lux room, separate and away from the testing room, for one hour prior to every assay. 70% ethanol was used to clean between each subject and trial in all described assays.

#### Open Field

Open field was used to detect locomotive activity and general explorative behavior, as previously described (Adhikari et al., 2018; Bales et al., 2014; Copping, Adhikari, Petkova, & Silverman, 2019; Copping et al., 2017). Individual mice were placed in a novel open arena for 30 minutes at 30 lux. Ethovision (Noldus) software was used to record sessions and analyze the total distance travelled during the 30 minutes and the average velocity of a subject during the session.

#### Beam-walking

To assess motor coordination and balance abilities, the beam-walking task was conducted. Subjects were placed, one by one, at one end of a 59-cm long beam, elevated 68 cm above a cushion, at the end of which a darkened goal box (12-cm diameter cylinder) was placed on the far end of the beam to provide motivation to walk across. On a first training day, three trials on a large diameter (35-mm) beam are conducted for mice to become accustomed to the task. Animals that had scores of 60 seconds for all three trials on the training day are excluded from analysis; no animals were excluded during this experiment. On the next day, animals were placed on three beams of different diameters (35-, 18-, and 13-mm) in order of least to most difficult (widest to narrowest). Two trials per beam are conducted with an inter-trial rest interval of at least 30 minutes. Each trial is a maximum of 60 seconds. Any falls off the beam were recorded as 60 seconds. The average latency to traverse the beam for the two trials was recorded.

#### Rotarod

An accelerating rotarod task was employed to further assess motor coordination and motor learning (Ugo Basile, Schwenksville, PA). Mice were placed on an accelerating cylinder that slowly accelerated from 4 to 40 revolutions per minute over a five-minute period. Latency to fall off the cylinder or no longer comply with the task (*i.e.,* cling to rod for more than two full rotations instead of walking forward) was recorded per trial with a maximum of 300 seconds. Mice tested for three consecutive days with three trials per day separated with an hour inter-trial interval to prevent fatigue and were tested for three consecutive days. Results from each day’s trials were averaged together, as the field’s standard (Yang et al., 2012).

#### Gait Analysis

The DigiGait imaging system was used to analyze gait (Mouse Specifics Inc, Boston, MA). DigiGait utilizes a ventral plane camera underneath a transparent treadmill belt that generates digital paw prints as a subject walk or runs (T. G. Hampton, Stasko, Kale, Amende, & Costa, 2004). Initial characterizations and validations were performed using protocols by (Amende et al., 2005; T. G. Hampton & Amende, 2010; T. G. Hampton et al., 2004). Protocols were adapted for AS by (Grieco, Gouelle, & Weeber, 2018) and personal communications with FAST and Dr. Jessica Duis.

The day after weaning, mice were habituated in the walking chamber for one minute and then the belt was turned on and slowly increased from 5 cm/s to the target belt speed of 20 cm/s. On all subsequent testing days, mice were placed in the walking chamber for one minute and the belt speed was set to 20 cm/s. Subjects unable to walk at the target speed for at least five seconds were allowed to rest and were retested. Subjects were recorded walking for 4-5 seconds, allowing for at least 10 full strides. Videos were taken when animals were running forward and were retaken if there were instances of subjects jumping, using the front bumper as an aid, walking diagonally across the camera view, or sliding backwards into back bumper.

Frames were digitized, and relevant gait parameters were analyzed using DigiGait analysis software as previously described (T. Hampton, 2004; T. G. Hampton et al., 2004). Quality control steps in analysis were conducted blind to genotype. Left and right fore- and hindlimbs were averaged together per subject for each metric.

### Statistical analysis

All statistical analysis was conducted using GraphPad PRISM software. Open field data was analyzed using Student’s t-test. Two-way repeated measures ANOVA was used to assess the beam walking, rotarod tasks, and adult DigiGait data with planned post-hoc multiple comparisons assessing performance between genotypes at each rod, day, and limb respectively. Longitudinal DigiGait data was analyzed using repeated measures mixed effects models with planned Sidak post-hoc multiple comparisons of genotype performance at each time point as well as multiple comparisons between all timepoints within genotype for each parameter studies. Data was analyzed for sex differences before pooling; there were no sex differences found in motor assays or gait metrics (**Supp. Fig. 5**). Additional, a full report of statistics is available in **Supplementary Table 1**.

## Results

### Global motor deficits

Adult AS mice exhibited global motor deficits as assessed by three gold-standard rodent behavioral assays at 4 months of age. As previously reported, AS mice showed reduced activity in a novel open arena exploration task (**Fig. 1A-B**). AS mice traveled less during the exploration period, traversing, on average, 25% less than WT (**Fig. 1A**, *p* = .003). Moreover, during the same period of time, AS mice showed reduced speeds, on average producing only 80% of the speed of WT mice (**Fig. 1B**, *p* = .026). In addition to gross locomotive ability, we assayed other motor abilities such as balance, coordination, and motor learning using the beam walking and accelerating rotarod tasks and saw that AS mice had impairments in both tasks (**Figs. 1C-D**). AS mice demonstrated characteristic low performance on accelerated rotarod on all three days (**Fig. 1C**_genotype_, p <.05). AS mice showed significantly increased latencies to cross more difficult beams, demonstrating an impairment in balance and coordination (**Fig. 1D**, Holm-Sidak posthoc analysis, AS vs. WT, rod 1: ns, rod 2: p =.0162, rod 3: p < .0001).

**Figure 1.**
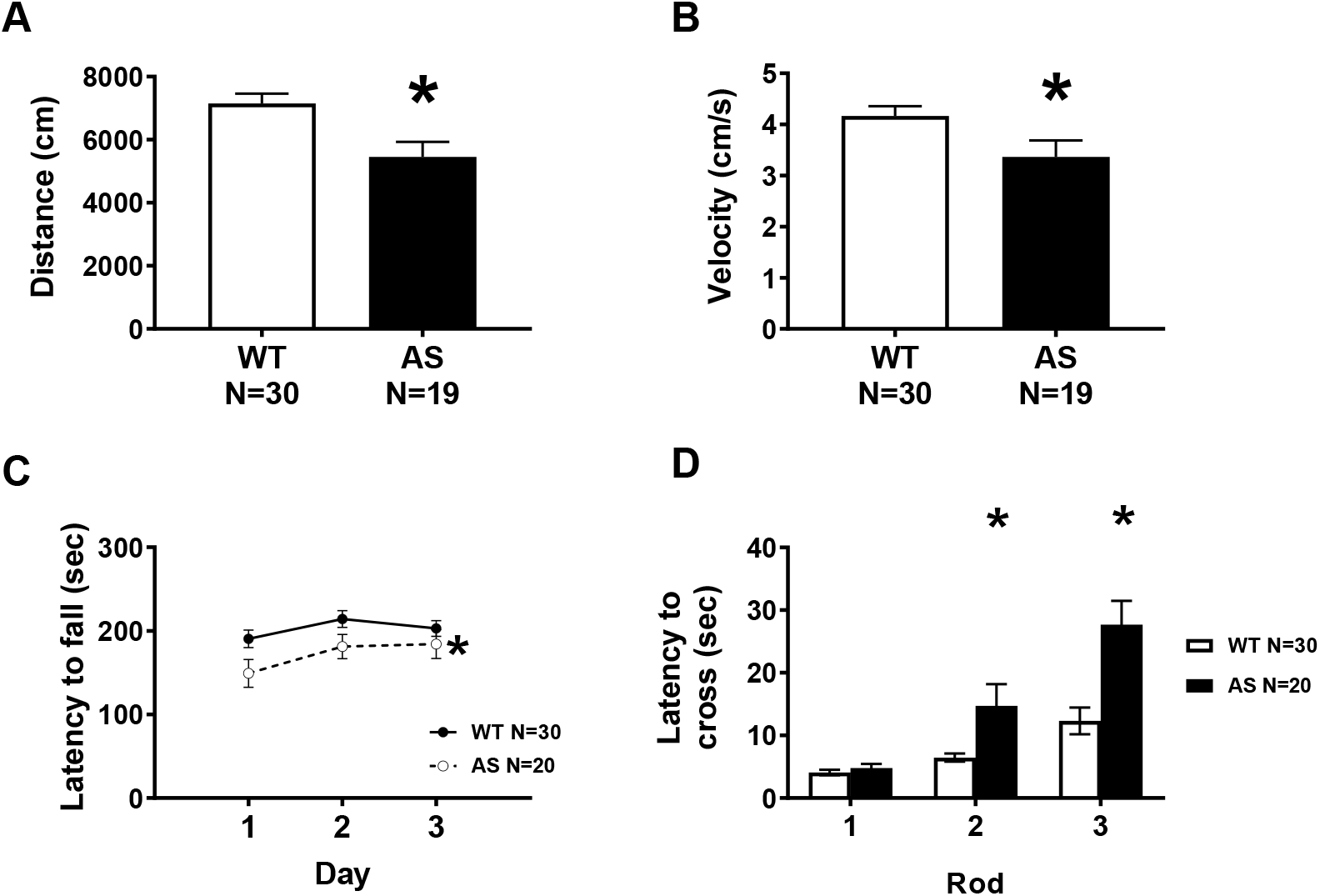
Adult AS mice exhibit severe motor deficits in locomotor exploration, speed, motor coordination, balance, and motor learning. **A-B.** Adult AS mice were given a 30-minute exploration period in a novel chamber. **A.** AS mice demonstrated low locomotive activity, as they traversed less distance, compared to wildtype mice. B. Moreover, during the exploration period, AS mice moved significantly more slowly than WT. C. In the accelerating rotarod, AS mice showed a significant decrease in latency to fall across all three days, suggesting an impairment in motor learning and coordination. **D.** Motor balance and coordination were assessed in a beam walking assay. Mice were trained to traverse a beam and their latency to cross the beam was recorded on progressively thinner and more difficult beams. AS mice took significantly longer than WT mice to cross the more difficult beams. * p <.05 in posthoc comparison between WT and AS. **(A-D)** Data represented as mean ±SEM. **(A-B)** * p<.05 Student’s *t*-Test between genotypes. **(C-D)** * p<.05 main effect of genotype in repeated measures two-way ANOVA.

### Adult gait

AS mice showed abnormal spatial and temporal subcomponents of gait compared to age and sex matched WT littermate controls. Spatial parameters included stance width, step length, and stepping frequency, also known as cadence. Stance width, defined as the distance between right and left limbs when both paws are on the ground, was statistically wider in the forelimbs and trending in the hindlimbs of AS mice, compared to WT (**Fig. 2A**, AS vs WT Holm-Sidak posthoc analysis, fore: p <.0001, hind: p =.07). Additionally, AS mice took longer steps, with an average 20% longer step or stride length in both the fore and hind limbs (**Fig. 2B**, AS vs WT Holm-Sidak posthoc analysis, fore: p <.0001, hind: p <.0001). Accordingly, stride frequency was decreased in AS fore and hind limbs (**Fig. 2B**, AS vs WT Holm-Sidak posthoc analysis, fore: p <.0001, hind: p <.0001). The abnormal wider and elongated gait adult AS mice may indicate lower ability during ambulation. Total stride duration and its four phases of temporal gait pattern were analyzed: swing, during which a paw has no contact with the ground, braking, during which a paw transitions from the swing phase and steps onto the ground into the stance phase, propulsion, during which a paw lifts off the ground and into the swing phase, and stance when the paw makes full contact with the ground. AS mice required more time to make one stride as the stride duration was significantly increased (**Fig. 3A**, AS vs WT Holm-Sidak posthoc analysis p, fore: p <.0001, hind: p <.0001). Within the stride, both stance and swing times were increased as AS mice spent more time with their paws on the ground and in the air compared to WT mice (**Fig. 2B-C**, both swing and stance AS vs WT Holm-Sidak posthoc analysis, fore: p <.0001, hind: p <.0001), which suggests the increase in stride time was not determined by a single component of the gait cycle. In the transitional phases of gait, we observed elevated propulsion time both the fore and hindlimbs in AS gait (**Fig. 3D**, AS vs WT Holm-Sidak posthoc analysis, fore: p <.0001, hind: p <.0001) whereas time required to decelerate and brake was significant only in the forelimbs (**Fig. 3E**, AS vs WT Holm-Sidak posthoc analysis, fore: p <.0001, hind: ns). We also looked at the percent of the stance phase where both paws are on the treadmill, referred to as % shared time and % double support. Organisms with poor balance tend to have increased double support as it increases stability; however, we observed that the AS mice actually had a decreased % shared time (**Fig. 2F**, t_(47)_ = 3.485, p =.0011). These phenotypes were replicated in a second, independent cohort (**Supp. Fig. 4).**

**Figure 2.**
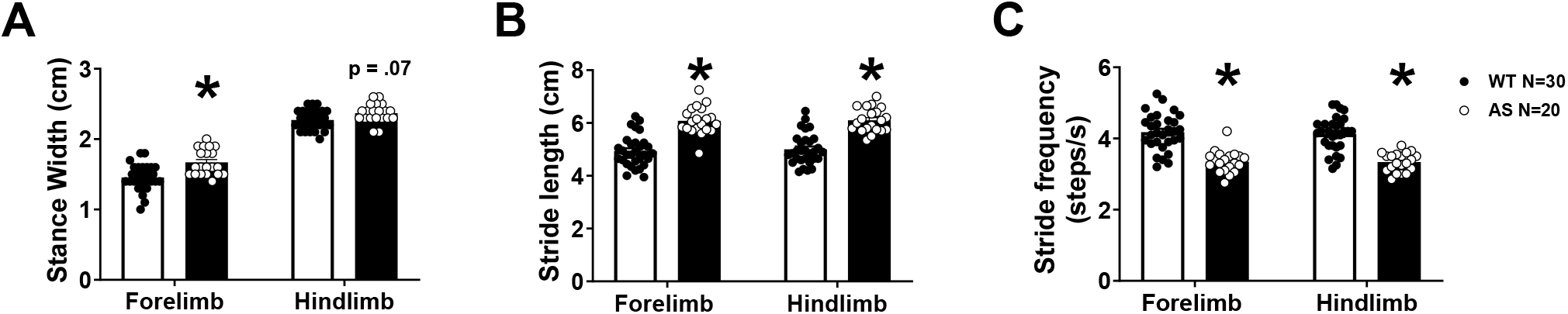
AS mice exhibit deficits in spatial metrics of gait compared to WT controls. **A.** During ambulation, stance width between left and right limbs during maximum stance was wider in AS mice in both fore and hindlimbs, potentially indicating an instable posture. **B.** AS mice took longer strides in both fore and hindlimbs and therefore, conversely, **C.** the step frequency, or cadence, was significantly reduced. Bars represent mean ± SEM * indicates p <.05, Student *t*-test between genotypes.

**Figure 3.**
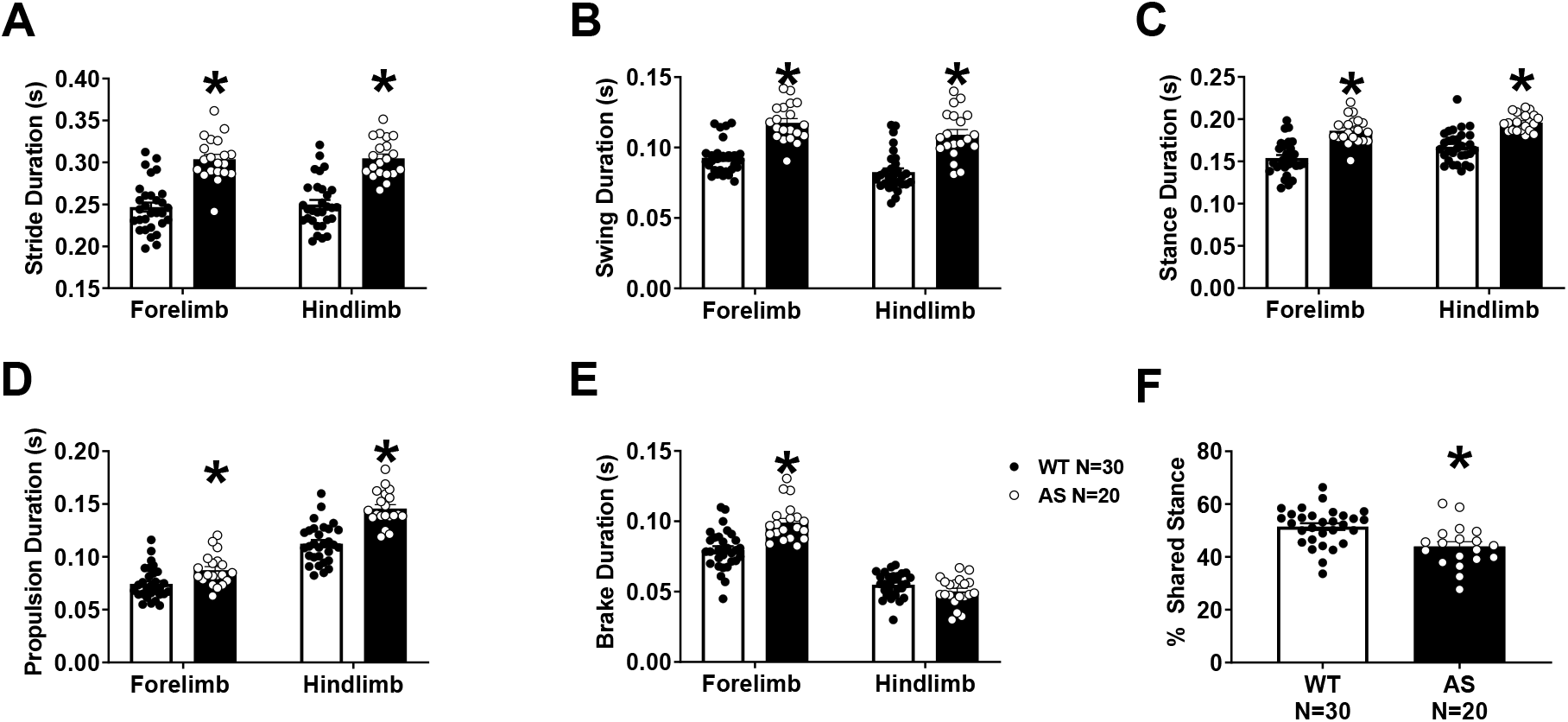
AS mice exhibit deficits in temporal metrics of gait compared to WT controls. **A.** Stride duration was elevated in both fore and hindlimbs of adult AS mice, compared to WT controls, requiring more time to make a single step. **B.** AS mice showed an increased duration spent in the swing phase, when the paw has no contact with the ground, compared to WT. **C**. Converse to swing, it was expected that the AS mice spent less time in the stance phase of the stride. **D.** Within the stance phase, propulsion is the lift-off of the paw from the ground into swing, propelling the animal forward. The AS mice show increased time spent in this phase. **E.** At the opposite end of the stance is braking, when the paw exits the swing and meets the ground, which was also shorter in AS, but only in the forelimbs. This is likely related to the specialized role of steering forelimbs are primarily used for in quadrupeds. **F.** Curiously, AS mice showed reduced time spent in % shared stance, also known as double support. Bars represent mean ± SEM * indicates p <.05 (A-E) between genotype post hoc comparison, (F) Student’s T-test.

### Gait across development

Next, we wanted to elucidate the progression of observable gait abnormalities in AS mice and whether we could observe an onset and specific developmental pattern of nuanced motor disabilities. We assessed gait from weaning until six months of age in AS mice to investigate the trajectory of the abnormal gait we observed in adulthood. Over this time course, AS mice showed reliably wider stance widths (**Fig. 4A**, fore: F_genotype (1,48)_ = 22.02, *p* < .0001, hind: F_genotype (1,48)_ = 11.65, *p* < .0001). As previously, we utilized Holm-Sidak multiple comparison tests to evaluate changes in gait parameters within genotypes, between time points. To that end, interestingly, while AS mice had wider stances, both AS and WT mice showed the same developmental pattern of stance width with steady increases over development and a sharp increase between PNDs 120 and 180. A longer step length was observed as early as the time of weaning in AS hindlimbs, and PND30 in forelimbs and it continued to increase throughout development, whereas WT mice showed a slight but significant increase in step length between PNDs 22 and 30 and then showed a stable, mature step length until after PND120 (**Fig. 4B**, fore: F_genotype (1,48)_ = 75.78, *p* < .0001, hind: F _genotype (1,48)_ = 76.48, *p* < .0001). As expected, we saw the converse in step frequency; AS had a decreased cadence that worsened throughout their lives, while WT mice showed only moderate changes between PNDs 22 and 30 and 120 and 180 (**Fig. 4C**, fore: F_genotype (1,48)_ = 63.75, *p* < .0001, hind: F _genotype (1,48)_ = 70.80, *p* < .0001). In humans, as gait matures from first steps into independent walking, the stance width between the feet decreases, step length increases as limb lengths also increase, and cadence, or step frequency, increases as a child’s speed increases. Here, speed is held constant therefore, while stride lengths increase as mice grow, the step frequency inversely decreases.

**Figure 4.**
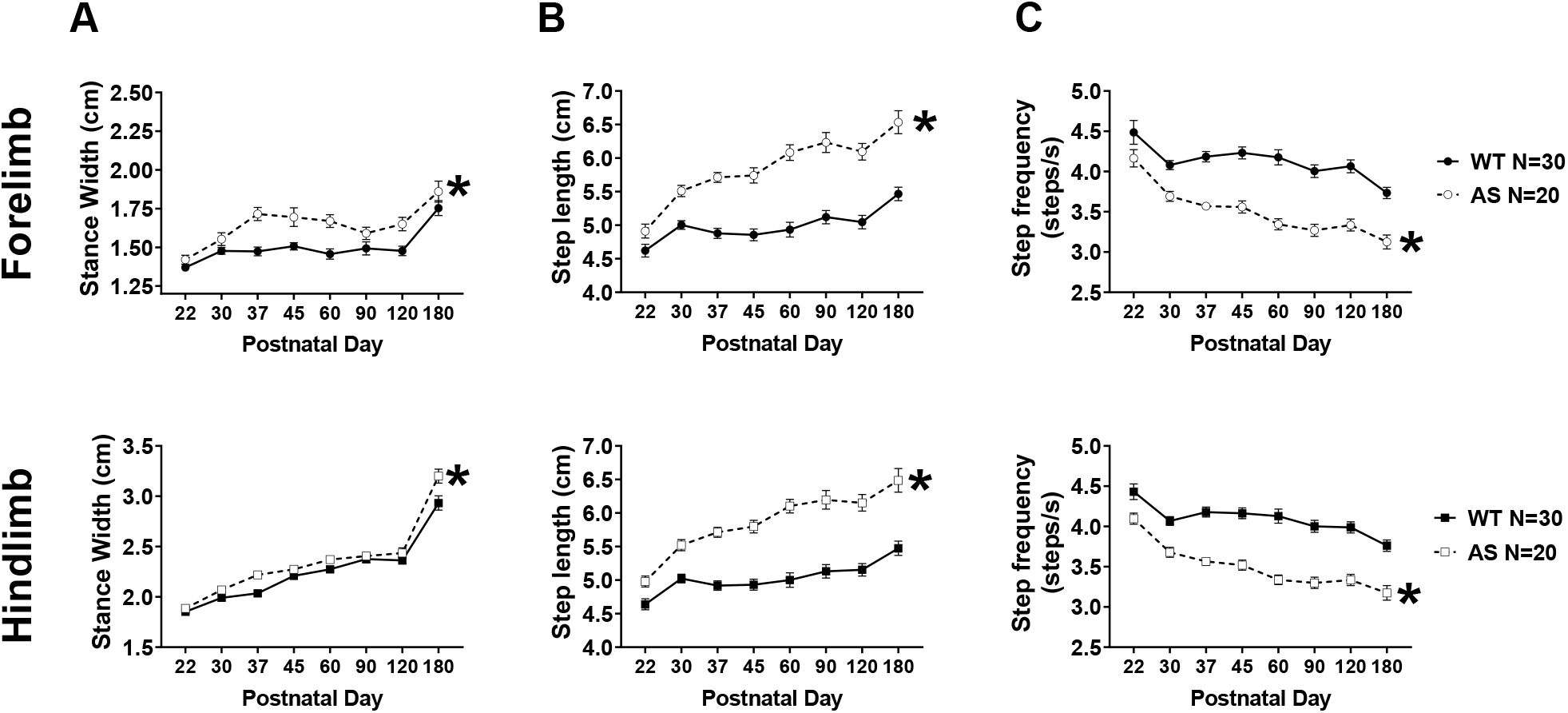
Across development, AS mice exhibited slower progression across all spatial gait indices. **A.** Stance width between right and left limbs in both fore (top) and hindlimbs (bottom) increased slightly throughout development but was wider in the AS mice, compared to WT mice, throughout. At the oldest time point, both WT and AS mice showed a sharp increase in stance width with a similar slope. **B.** Similarly, AS mice exhibited increased step lengths throughout the experiment and the discrepancy in step length was present at weaning at PND22. Additionally, while WT mice had a stabilized step length after PND30, step length continued to increase in AS mice and began stabilizing around after PND60. The same pattern was observed in both fore and hindlimbs **C.** Expectedly, the stepping frequency showed the converse: in both limbs, AS mice showed reduced step frequency that was present at weaning and progressed throughout development, unlike the WT mice which showed a relatively stable, mature cadence after PND30. * p <.05, main effect of group of two-way repeated measured mixed effects model.

When we looked at temporal subcomponents of gait, we saw that AS mice had disrupted developmental gait trajectories, with complex temporal patterns. Across all the temporal parameters reported, we observed that AS mice displayed a significantly elongated stride time in both the fore and limbs that was continuously prolonged while WT stride time stabilized after PND30 and until PND120, at which time both groups showed an age-related increase in stride time (**Fig. 5A**, fore: F_genotype (1,48)_ = 75.92, *p* < .0001, hind: F_genotype (1,48)_ = 76.54, *p* < .0001). AS mice spent more time in the stance phase than WT mice with both groups showing similar developmental patterns (**Fig. 5B**, fore: F_genotype (1,48)_ = 55.22, *p* < .0001, hind: F_genotype (1,48)_ = 55.03, *p* < .0001). AS swing time was also detectably longer in the forelimbs at PND30, but this aberration was present at weaning in the hindlimbs (**Fig. 5C**, fore: F _genotype (1,48)_ = 84.17, *p* < .0001, hind: F _genotype (1,48)_ = 64.94, *p* < .0001). Corroborating our earlier observations, we detected a developmental pattern: a stabilization of gait metrics in WT mice after PND30 and an age-related increase between PND120 and 180 that was also observed in AS mice. AS mice spent more time in the propulsion phase of gait compared to WT, with both groups showing the same pattern across time (**Fig. 5D**, fore: F_genotype (1,48)_ = 34.07, *p* < .0001, hind: F_genotype (1,48)_ = 58.53, *p* < .0001). AS mice did not show a deviation from WT in braking of the hindlimbs, but did show increased braking time in the forelimbs (**Fig. 5E**, fore: F _genotype (1,48)_ = 29.31, *p* < .0001, hind: F_genotype (1,48)_ = 4.599, *p* = .0371). The pattern of braking duration across time was unique compared to the other temporal metrics. A steep increase in braking time between PND22 and 30 was revealed in both groups; WT had a stabilization until PND120, while the AS mice continued to spent more time in the brake phase until PND60, after which point the brake time decreased and was not significantly longer than WT. At PND60, AS mice showed significantly reduced time in double support when both paws are simultaneously on the ground (**Fig. 2F**), we saw that this was not the case across the developmental time window we studied (**Fig. 5F**). There was a significant difference between the genotypes (F_genotype (1,48)_ = 3.485, *p* = .0011), although Holm-Sidak multiple comparisons could not detect that double support was not different from WT at early time points of PND 22 and 30 and later time points at PND120 and 180, so while WT mice showed an increase in double support early in life, AS showed slow increases and more drastic increases between advanced time points.

**Figure 5.**
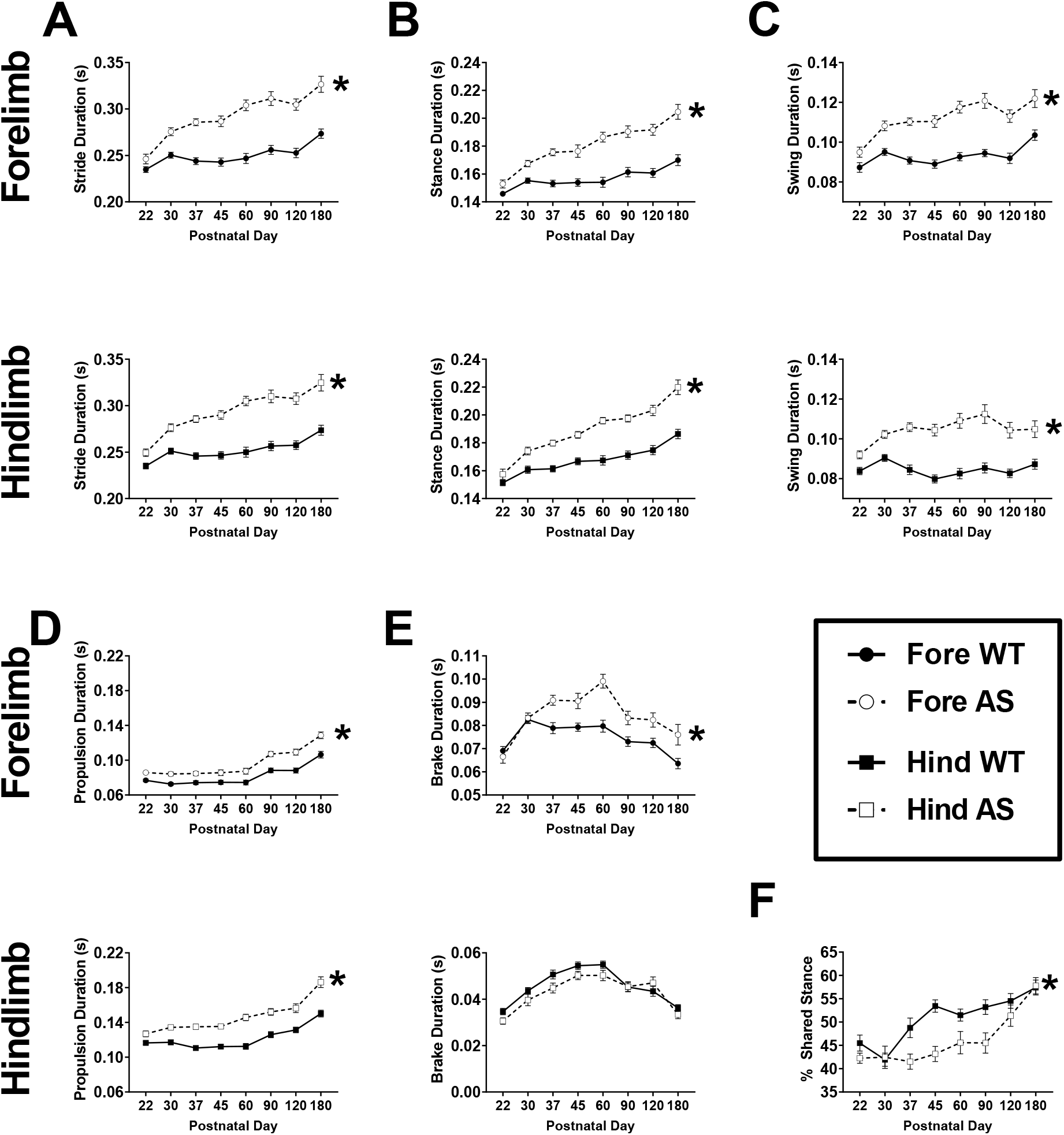
AS mice exhibited aberrant temporal metrics of gait beginning at PND30 with progressive worsening over time. Temporal subcomponents of gait were detectably different in AS mice as early as weaning, but typically at PND30, and continued to progress while WT mice showed maturation and stabilization of gait parameters. **A.** Stride duration was significantly higher in the fore and hindlimbs of AS mice, compared to WT. **B.** Aberrant stance time was observed in AS mice, in both limbs. **C.** Similar to stance time, swing time was also elevated in AS mice. Interestingly, elevated hindlimb swing time was detected earlier at PND22. **D.** Propulsion time was increased in the AS group PND30 onwards, which suggests more time is required to produce force necessary to initiate the next step. **E.** Time spent in the braking phase was only significantly higher in the forelimbs of AS mice, likely due to the specific use of forelimbs in steering. This difference emerged at PND37. **F.** Double support, or % shared stance time, in the hindlimbs, was significantly different between AS and WT mice over the lifespan. * p <.05, main effect of group of two-way repeated measured mixed-effects model.

We observed that many abnormalities in AS gait present as early as the first time point, weaning. Mice, of either genotype, were unable to walk consistently and continuously long enough at the requisite speed to capture high quality videos for analysis earlier than PND22. We attempted to observe whether ambulation abnormalities arose earlier by performing a neonatal circle transverse task and saw that AS pups gained proficient crawling and walking skills later than WT, suggesting motor deficits are present early in life and progress (**Supp. Fig. 2**).

## Discussion

This is the first report of precisely quantified spatiotemporal metrics of gait and motor patterns across development in AS model mice, a uniquely translational metric, over development. Motor deficits have been key in the study of rodent models of AS, as dysfunction on the rotarod and reduced activity have been the most consistently reported motor behavioral phenotypes, likely related to cerebellar dysfunction (Berg et al., 2020; Born et al., 2017; Cheron, Servais, Wagstaff, & Dan, 2005; Dodge et al., 2020; Huang et al., 2013; Sonzogni et al., 2018). These have been and continue to be key reproducible observations in AS. Our report extends this work and highlights: 1) global motor deficits in AS mice, 2) substantial gait aberrations in adult AS mice, 3) evidence that gait abnormalities are detectable early in life and progressively decline throughout aging.

We corroborated earlier work that demonstrates a severe global motor deficit in AS mice. Importantly, we discovered numerous aberrant quantitative gait parameters that indicate an instable and weakened gait, similar to clinic reports by Dr. Jessica Duis (personal communication) and (Grieco et al., 2018). We observed wider stances in both limbs in adults and throughout development, suggesting instable gait as stance widening serves as a compensatory measure for ambulation. Spatial gait was further disrupted as AS mice took longer, fewer, steps compared to WT. This may be less related to stability but capability of AS animals; one conjecture is that the speed chosen represents a walk for motorically normal WT mice but may represent more of an effortful run for AS mice, requiring a gait pattern closer to running which is characterized by longer strides and reduced stride frequency. At a higher target speed of 36 cm/s, when both animals are running, stride lengths were longer, compared to slower speeds, and there were fewer differences in gait patterns between genotypes (**Supp. Fig. 6**). The reduced double support observed in AS mice as adults and across much of the developmental timeline may also support this hypothesis as double support typically decreases as a function of speed in humans and quadrupeds. Temporal indices of gait were also altered in AS mice, by lengthened phases of the gait cycle and of the cycle as a whole. Overall stride time was increased but not as a function of any singular component; both swing when a paw is in the air, and stance, when it is on the ground, times were increased in both sets of limbs. This slowness may be linked to the longer steps taken in AS subjects to create an abnormal pattern of slower and longer strides in order to maintain the requisite treadmill speed. Of particular importance was the increased propulsion in AS mice. Propulsion time is the time a paw requires to transition from the stance into the swing phase, effectively pushing off the ground and propelling the animal forward. We discovered elevated propulsion time in AS gait which, aligned with the increased paw drag AS mice (**Supp. Fig. 1**), suggests a potential limb weakness not detectable in other less nuanced standard motor behavioral assays. AS subjects required more time to generate the same force to propel themselves into the next step compared to WT because they lacked the equivalent strength. Additionally, we saw limb-specific patterns in the braking time where AS mice had altered braking time in the forelimbs but not the hindlimbs. This is likely due to the separate roles of fore and hindlimbs in quadrupeds; hindlimbs are primarily used to push off and propel the animal forward while the forelimbs are preferentially used for steering and controlling deceleration. Combined together, the augmented braking and propulsion suggest that AS mice may either have lack of motor control as they rapidly place a paw onto the ground or reduced muscular strength, as they require more time to produce the same force necessary to propel into the next step. A limitation of the current work is that such detailed and nuanced motor deficits have yet to be commonly identified or reported in preclinical models of NDDs. Our goal is to continue this examination and provide this data in a variety of preclinical models as it may predict go/no-go clinical decisions in the future.

We detected that many abnormal gait metrics identified in adulthood were present at earlier ages. Weaning was chosen as a first time point because mice younger than PND21 were not able to successfully walk at the requisite speed of 20 cm/s that has been established as the standard speed of data collection. A secondary limitation was that we could not explore many slower speeds at the present time. We are working to identify how the DigiGait can be modified for earlier developmental timepoints. Differences in gait phenotypes were significant and robust by postnatal day 30, and in many cases, sooner. This suggests a general delay or an underdevelopment of the motor skills required for ambulation. To test this, we tracked ambulatory skills during the neonatal period and did see a delay in successful ambulation but were unable to delineate data’s underpinning (**Supp. Fig. 2**). Longitudinal assessments are imperative for several reasons and gait proves a reliable, translational assay to accomplish this goal unlike some other behavioral assays. For one, ambulation in isolation requires a novel arena and has test-retest effects (i.e. signal is lower with each exposure). Rotarod does not accommodate for confounding contributions such as background strain and weight and also has test-retest effects (i.e. signal is changes with exposure). Moreover, to our knowledge, there is no known neural circuitry of human rotarod in existence. DigiGait avoids many of these pitfalls and has been to be a versatile tool, sensitive to various treatments, and has a record of use as a strong longitudinal/repeat testing outcome measure (Adhikari et al., 2021; Akula et al. 2020; Hansen & Pulst, 2013; Orfany et al., 2020; Zhan et al., 2019). Treadmill gait analysis using ventral imaging, such as DigiGait and the CatWalk, allow for unhindered access to the ambulating mouse to detect more nuanced and fine-grained phenotypes (Lei et al., 2014). Assays with increased sensitivty of mouse motor ataxia may provide greater understanding of the pathological profile and could be used in coorboration with standard assays.

A gap in preclinical research is a lack of focus on onset and progression of NDD relevant symptoms, as essentially all preclinical behavior is performed in adults, foregoing identifying earlier onsets of detectable behavioral differences and tracking progression, be it improvement or decline, over time. Furthermore, a lack of consensus in literature on the appropriate age for behavioral assessment, following therapeutic interventions, may confound obtaining true positive, reliable, and reproducible results. Testing different ages at which times the degree of impairment is not ascertained and well-known can lead to both false positive (type I) and false negative (type II) errors.

Motor impairment has been studied in mouse models using the accelerating rotarod and exploration of an open field, but these assessments lack rigor, rely on non-analogous mouse circuitry, have mouse-specific confounds such as weight, age, and background strain, and are not translational metrics (Mao, Ogata, Ora, Tanaka, & Miyake, 2021; Shiotsuki et al., 2010; Shoji & Miyakawa, 2021; Stroobants, Gantois, Pooters, & D’Hooge, 2013). Many pharmaceutical clinical trials have “failed” to move forward successfully despite preclinical “rescues”; our work suggests it is possible that tasks lacking translational specificity and analogous neural circuitry contribute to this phenonmenon (Gordon, 2019); in other words, the weaknesses of the outcome measures preclinically are equally as vital as the lack of quantifiable outcome measures clinically. We predict this will reduce the “valley of death” or “lost in translation” gap in bench to bedside research. An example is the latency to fall off the rotarod which is confounded by weight, age, mice background, repeated testing, and lacks complexity in its behavior (Deacon, 2013). This has also been true for other models of NDDs where underdeveloped and insensitive outcome measures have resulted in failure to show meaningful change in potentially promising drug studies (Erickson et al., 2017). Automated gait analysis may overcome many limitations, by presenting a repeatable, robust, reproducible quantitative method of assessing multiple metrics and indices and producing analogous, if not identical, components to clinical gait analysis. Indices and metrics congruent to human gait studies should be more readily included in future behavioral phenotyping deep dives (Akula et al., 2020; Hampton et al., 2004; Hansen & Pulst, 2013; Lei et al., 2019; Lei et al., 2016).

Our study showcases translational and longitudinal use of quantitative motor metrics for a genetic NDD. Clinically, quantitative gait research has gained traction across many NDDs and the idea to tailor attention on gait in preclinical studies is now also gaining momentum (IDDRC working group, AGENDA working group, personal communications) for Rett syndrome, neurofibromatosis, Down syndrome, and other NDDs (Hampton et al., 2004; Jequier Gygax et al., 2020; Kennedy et al., 2020; Layne et al. 2018; Rahn et al., 2020; Summers et al., 2015; Wilson et al., 2020). Data from our work showed analogous measures of gait phenotypes in rodents that are being quantified currently in AS clinics (Jessica Duis MD, personal communication). Of impact is that these metrics are not specific to any single syndrome but represent measures that can broadly be applied across many different NDDs with impacted motor development and gait patterning. We identified a series of parameters that demonstrate maladaptive gait progression, potentially linked to reduced limb strength and coordination. We plan to expand this work to other NDDs, such as Phelan-McDermid Syndrome, Duplication15q Syndrome, and genetically identified syndromic autism spectrum disorders, for which there is a growing body of evidence that there exists a relationship between degree of motor deficit and other behavioral domains, such as social communication and cognition (Copping et al., 2016; DiStefano et al., 2016; Finucane et al., 2016; Shumway et al., 2011; Soorya, Leon, Trelles, & Thurm, 2018). In summary, gait as an outcome measure is a convincingly robust, reliable, and consistent phenotype that showed strong translation in AS.

## Acknowledgements

This work was supported by generous funding from the Foundation for Angelman Syndrome Therapeutics (FAST) and the MIND Institute’s Intellectual and Developmental Disabilties Research Center (IDDRC; P50HD103256; PI LA).

