## Supplemental Figures for "Gait as a Quantitative Translational Outcome Measure in Angelman Syndrome"

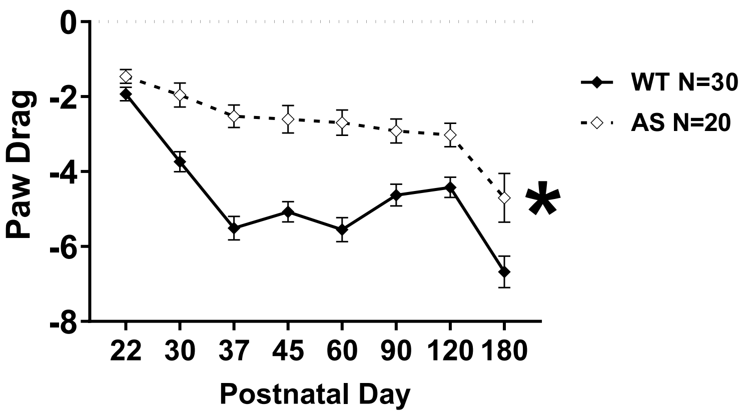


**Supplemental Figure 1: AS mice exhibited elevated paw drag, inferring poor hindlimb propulsion, compared to WT controls.**

Paw drag, calculated as the area under the curve during the propulsion phase, is elevated in the AS mice. There was a stark and significant deficit beginning at PND30. AS mice have less propulsive lift off. * $p < .05$ main effect of genotype in two-way repeated-measures ANOVA.

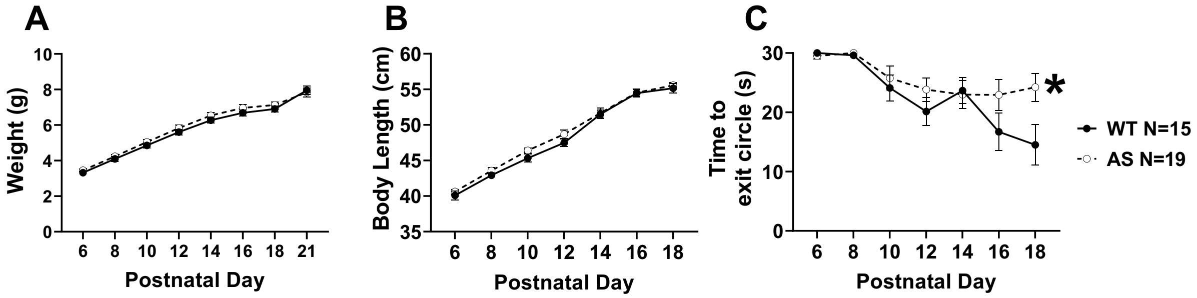


**Supplemental Figure 2: Gait development is delayed in AS mice.** Neonatal assessment showed that AS mice grew similarly to WT mice by weight (**A**) and length (**B**) but showed a delay in learning to walk. (**C**) Mouse pups typically begin crawling between PND7-11 and are fully ambulatory by PND20, at weaning Every other day between PND6 and 18, pups were placed in a small diameter circle and time required to move out of it was recorded as a proxy metric for development of ambulatory skills. AS mice showed delayed development of ambulatory skills as they took significantly more time to exit the circle, beginning at PND16. Gait abnormalities observed later using automated gait analysis appear to originate in neonatal development. * $p < .05$ main effect of genotype in two-way repeated-measures ANOVA.

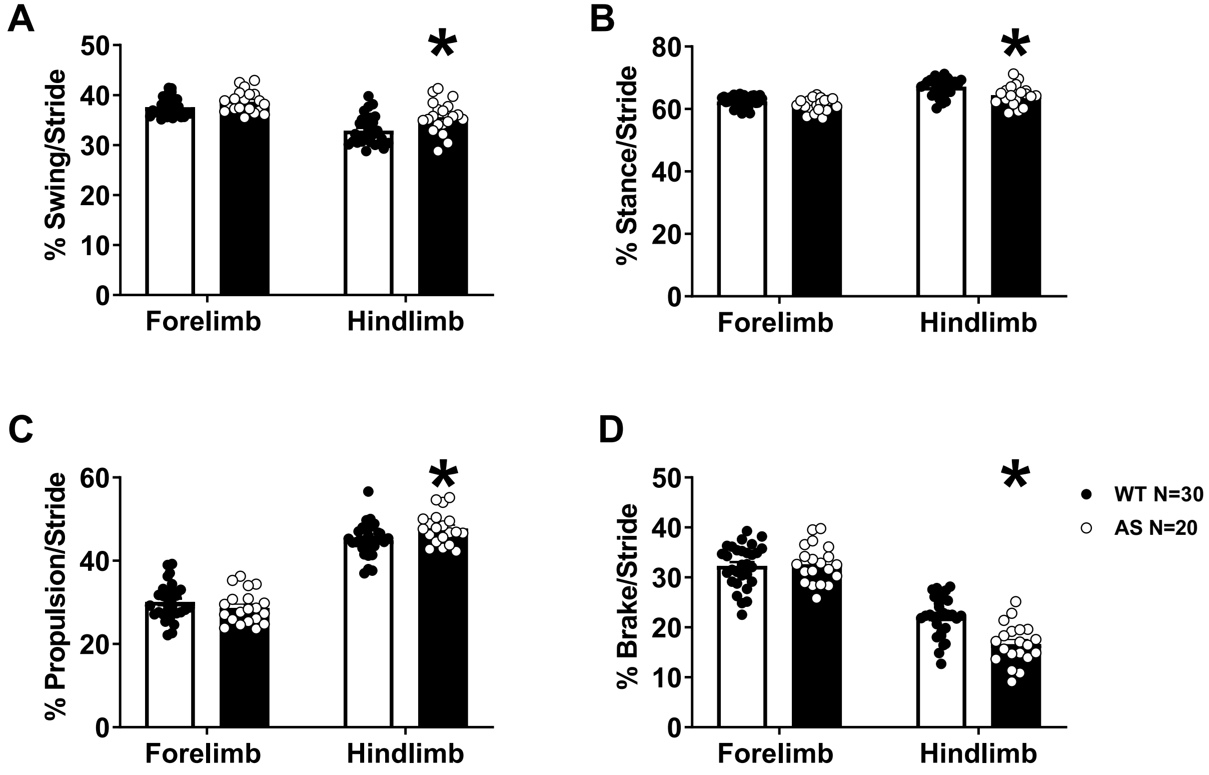


**Supplemental Figure 4: PND60 AS mice show abnormal portion of stride in temporal subcomponent gait metrics in hindlimbs.** We analyzed temporal subcomponents of gait as a function of the overall stride time. **A.** Hindlimbs of AS mice spent a greater portion of stride in the swing phase compared to WT while there was no difference in forelimbs. **B.** As stance and swing are converses, there was a decrease in % of stride spent in stance in hindlimbs. AS mice spend more time with their hindlimbs in the air, compared to on the ground. **C.** AS mouse hindlimbs spent a greater portion of the stride in propulsion, compared to WT mice. **D.** AS mouse hindlimbs spent a significantly smaller portion of stride in brake phase, compared to WT mice. Bars represent mean ± SEM. * indicates $p < .05$ between genotypes post hoc comparison

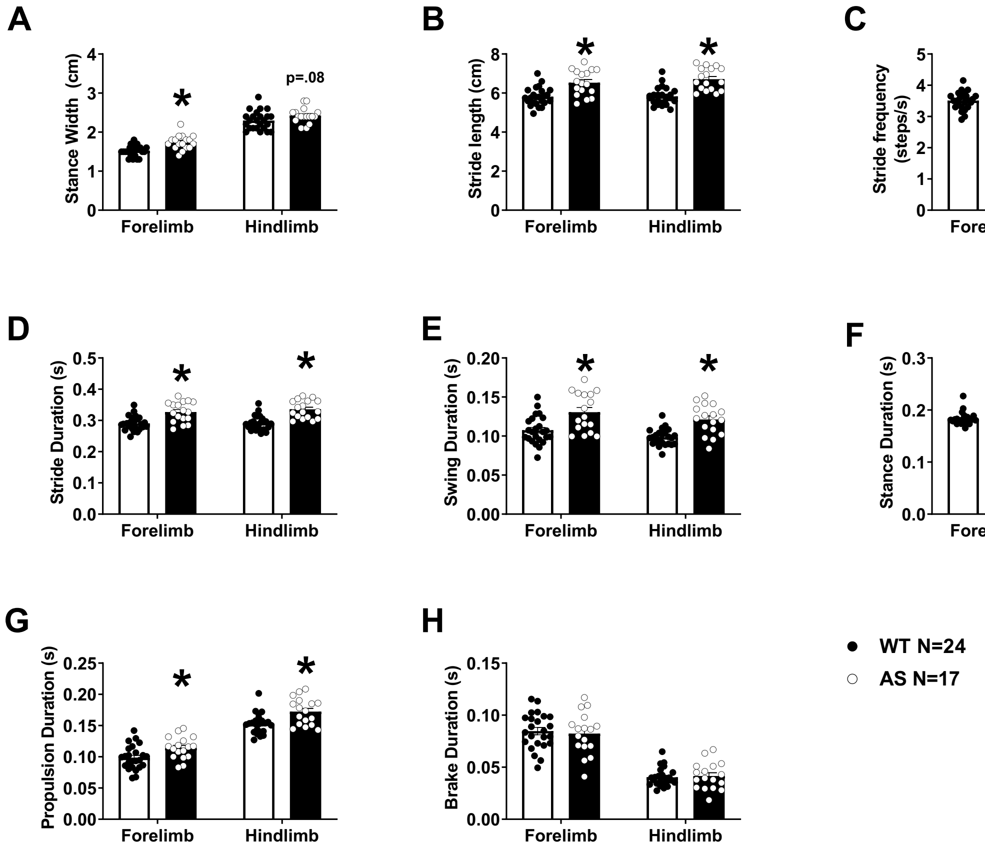


**Supplemental Figure 4: Gait aberrations in AS mice replicated in independent cohort.** An independent cohort was run through DigiGait at adulthood to demonstrate reproducibility of gait phenotype in adult AS mice. **A.** AS mice had significantly wider stance in the forelimbs and near significant in the hindlimbs, compared to wildtype controls. **B.** AS mice had elongated step lengths in both limbs and conversely showed significantly decreased stride frequency compared to WT controls (**C.**). **D.** Stride time was consistently increased in AS mouse fore and hindlimbs. **E.** Swing time when a paw is fully in the air was higher in both fore and hindlimbs of AS mice. **F.** The opposing phase, stance, when a paw is fully on the ground, was also increased in AS mice. **G.** AS mice took more time to push from stance into swing as propulsion was significantly increased in both sets of limbs, while **H.** brake time, the transition between swing into stance, was not altered in AS mice. Bars represent mean ± SEM. * indicates $p < .05$ between genotypes post hoc comparison

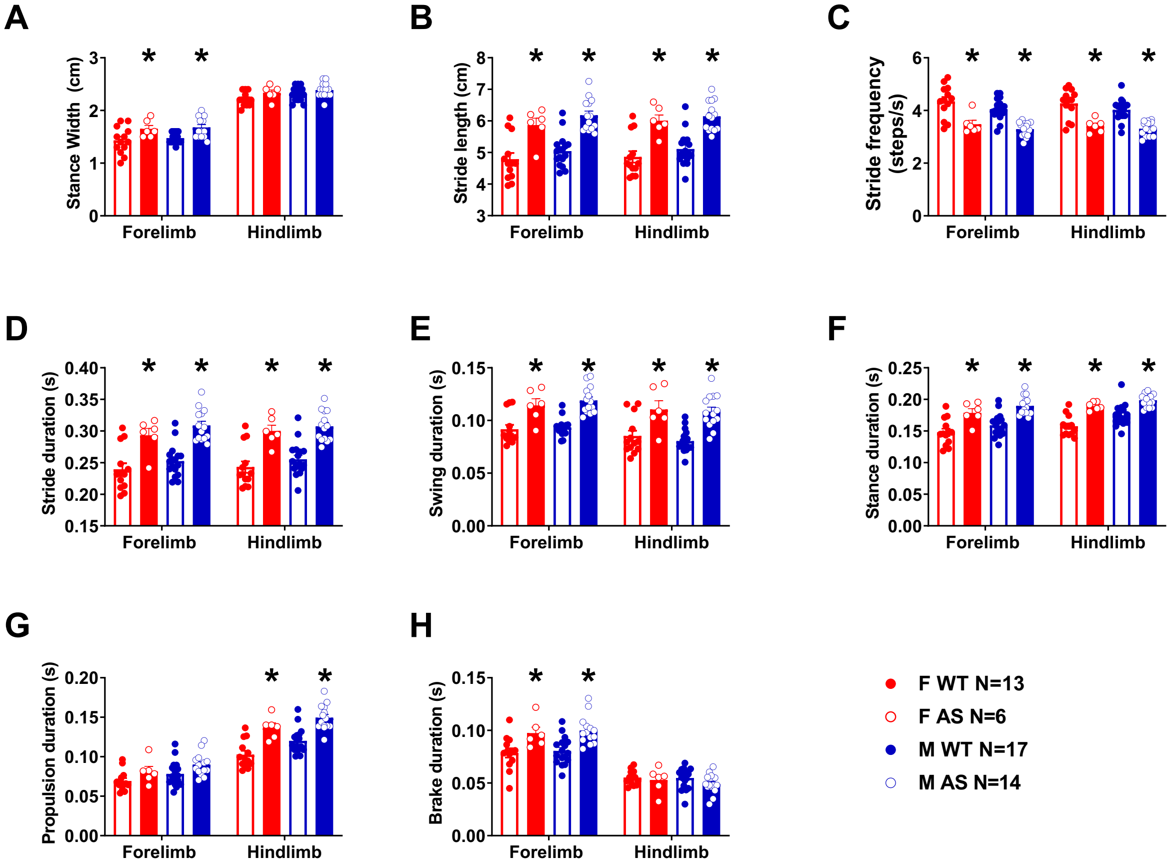


**Supplemental Figure 5: No sex differences observed in gait metrics in either WT or AS mice.** Gait metrics were analyzed by sex within genotype at each time point in the developmental study. For exemplar, data from PND60 mice was separated by sex to analyze for sex differences. Across all parameters, there were no sex differences between males and females within genotype; this remained true for all other PNDs included in the study and therefore data from both sexes was pooled together. **A.** Stance width was increased in forelimbs for both sexes of AS mice, but not the hindlimbs. **B.** Step length was elongated in both sexes and in both sets of limbs of AS mice. **C.** Conversely, stride frequency was decreased in both sexes and both sets of limbs. **D-F.** Overall stride duration as well as subcomponents of swing (**E.**) and stance (**F.**) were increased in male and female AS mice. **G.** AS mice of both sexes showed increased propulsion time in the hindlimbs and **H.** increased braking time in the forelimbs. Bars represent mean ± SEM. * indicates $p < .05$ within sex and limb, between genotypes post hoc comparison.

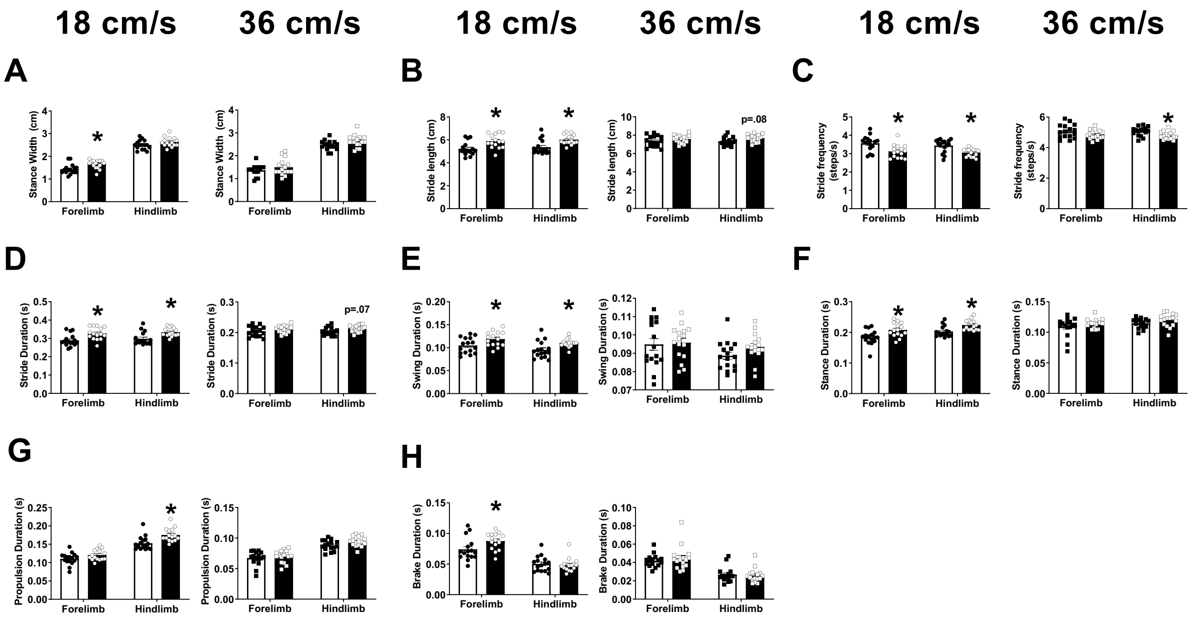


**Supplemental Figure 6: Gait aberrations in AS mice are consistent at slower speed but not at higher, running, speed.** We chose 20 cm/s as our target speed based on previous literature reports in mice using DigiGait but also investigated AS and WT performance in an independent cohort at two additional speeds based on literature (Hampton et al. 2004): 18 cm/s (left in panels) and 36 cm/s (right in panels) representing a slower walk and a high speed run respectively. **A.** AS mice showed increase stance width in the forelimbs at lower speeds but there was no difference in stance width at the higher running speed. **B.** AS mice showed longer strides in both limbs at 18 cm/s but only showed a trend towards increased step length in the hindlimbs at the fast 36 cm/s speed. **C.** Therefore, AS mice showed decrease stride frequency at the slower speed and decreased in the hindlimbs at the higher speed. **D.** Overall time to make a single stride was likewise increased in both limbs at 18 cm/s but only trended to an increase in hindlimbs when running at higher speeds. **E.** Swing time was increased in both sets of limbs of AS mice at 18 cm/s but not different from WT at the higher speed; the same pattern was observed in swing’s opposite, stance time (**F.**) **G.** Propulsion time was increased in hindlimbs when AS mice were walking at 18 cm/s but was not different from WT in either limb set at higher speeds. **H.** AS mice took more time to brake in the forelimbs at the slower speed while no differences were observed at 36 cm/s.
