## Supplemental Table 1 for "Gait as a Quantitative Translational Outcome Measure in Angelman Syndrome"

| **Test** | **# of Animals** | **Metric** | **Statistical Test** | **Statistic** | **p value** | **post hoc test** |  | **p value** |
| --- | --- | --- | --- | --- | --- | --- | --- | --- |
| Open Field | ***WT***  N=30  ***AS***  N=19 | Distance Travelled | Unpaired Two-Tailed  T-Test | T (47) = 3.094 | **p=.0033*** |  |  |  |
|  |  | Velocity | Unpaired Two-Tailed  T-Test | T (47) = 2.296 | **p=.0262*** |  |  |  |
| Accelerating Rotarod |  | Latency to Fall | Two Way Repeated Measures ANOVA | Genotype F (1,48) = 3.78  Day F (2,96) = 8.719  Interaction F (2,96) = 1.269 | **p=.05***  **p =.0003***  p =.2857 | ***WT*** vs  ***AS*** | Day 1 | p = .0671† |
|  |  |  |  |  |  |  | Day 2 | p= .1918 |
|  |  |  |  |  |  |  | Day 3 | p = .6613 |
| Beam Walking |  | Latency to Cross | Two Way Repeated Measures ANOVA | Genotype F (1,46) = 17.47  Rod F (2,92) = 34.57  Interaction F (2,92) = 7.56 | **p =.0001***  **p <.0001***  **p =.0009*** |  | Rod 1 | p< .9999 |
|  |  |  |  |  |  |  | Rod 2 | **p=.0162** |
|  |  |  |  |  |  |  | Rod 3 | **p<.0001*** |

| **Test** | **# of Animals** | **Metric** | **Statistical Test** | **Statistic** | **p value** | **Sidak Multiple**  **Comparison** | | **p value** |
| --- | --- | --- | --- | --- | --- | --- | --- | --- |
| Adult Digigait | ***WT***  N=30  ***AS***  N=20 | Stance Width | Two Way Repeated Measures ANOVA | Genotype F (1,48) = 17.13  Limb F (1,48) = 787.9  Interaction F (1,48) = 4.662 | **p =.0001***  **p <.0001***  **p =.0359*** | ***WT*** vs  ***AS*** | Forelimb | **p<.0001*** |
|  |  |  |  |  |  |  | Hindlimb | p = .0765† |
|  |  | Step Length | Two Way Repeated Measures ANOVA | Genotype F (1,48) = 50.90  Limb F (1,48) = 2.901  Interaction F (1,48) = 0.8685 | **p =.0001***  p =.095†  p =.3560 |  | Forelimb | **p<.0001*** |
|  |  |  |  |  |  |  | Hindlimb | **p<.0001*** |
|  |  | Step Frequency | Two Way Repeated Measures ANOVA | Genotype F (1,48) = 44.81  Limb F (1,48) = 1.612  Interaction F (1,48) = 0.8623 | **p <.0001***  p =.2103  p =.3577 |  | Forelimb | **p<.0001*** |
|  |  |  |  |  |  |  | Hindlimb | **p<.0001*** |
|  |  | Stride Duration | Two Way Repeated Measures ANOVA | Genotype F (1,48) = 50.65  Limb F (1,48) = 2.72  Interaction F (1,48) = 0.6907 | **p <.0001***  p = .1056  p =.410 |  | Forelimb | **p<.0001*** |
|  |  |  |  |  |  |  | Hindlimb | **p<.0001*** |
|  |  | Swing Duration | Two Way Repeated Measures ANOVA | Genotype F (1,48) = 45.17  Limb F (1,48) = 78.65  Interaction F (1,48) = 0.5844 | **p <.0001***  **p <.0001***  p =.4483 |  | Forelimb | **p<.0001*** |
|  |  |  |  |  |  |  | Hindlimb | **p<.0001*** |
|  |  | Stance Duration | Two Way Repeated Measures ANOVA | Genotype F (1,48) = 41.36  Limb F (1,48) = 66.87  Interaction F (1,48) = 1.956 | **p <.0001***  **p <.0001***  p =.1683 |  | Forelimb | **p<.0001*** |
|  |  |  |  |  |  |  | Hindlimb | **p<.0001*** |
|  |  | Propulsion Duration | Two Way Repeated Measures ANOVA | Genotype F (1,48) = 28.87  Limb F (1,48) = 628.5  Interaction F (1,48) = 27.71 | **p <.0001***  **p <.0001***  **p <.0001*** |  | Forelimb | **p = .0139*** |
|  |  |  |  |  |  |  | Hindlimb | **p<.0001*** |
|  |  | Brake Duration | Two Way Repeated Measures ANOVA | Genotype F (1,48) = 8.192  Limb F (1,48) = 284.5  Interaction F (1,48) = 30.29 | **p = .0062***  **p <.0001***  **p <.0001*** |  | Forelimb | **p<.0001*** |
|  |  |  |  |  |  |  | Hindlimb | p = .3136 |
|  |  | %Swing/  Stride | Two Way Repeated Measures ANOVA | Genotype F (1,48) = 9.488  Limb F (1,48) = 92.77  Interaction F (1,48) = 4.156 | **p=.0034***  **p <.0001***  **p =.0470*** |  | Forelimb | p=.2830 |
|  |  |  |  |  |  |  | Hindlimb | **p=.0007*** |
|  |  | %Stance/  Stride | Two Way Repeated Measures ANOVA | Genotype F (1,48) = 9.488  Limb F (1,48) = 92.77  Interaction F (1,48) = 4.156 | **p=.0034***  **p <.0001***  **p =.0470*** |  | Forelimb | p=.2830 |
|  |  |  |  |  |  |  | Hindlimb | **p=.0007*** |
|  |  | %Propulsion/  Stride | Two Way Repeated Measures ANOVA | Genotype F (1,48) = 0.5212  Limb F (1,48) = 723.5  Interaction F (1,48) = 11.50 | p = .4739  **p <.0001***  **p =.0014*** |  | Forelimb | p = .3962 |
|  |  |  |  |  |  |  | Hindlimb | **p=.0327*** |
|  |  | %Brake/  Stride | Two Way Repeated Measures ANOVA | Genotype F (1,48) = 7.298  Limb F (1,48) = 387.0  Interaction F (1,48) = 19.6 | **p = .0095***  **p <.0001***  **p <.0001*** |  | Forelimb | p = .9424 |
|  |  |  |  |  |  |  | Hindlimb | **p<.0001*** |
|  |  | % Shared Stance | Unpaired Two-Tailed  T-Test | T (47) = 3.485 | **p=.0011*** |  |  |  |
|  |  | Paw Drag | Unpaired Two-Tailed  T-Test | T (48) = 5.985 | **p <.0001*** |  |  |  |

| **Test** | **# of Animals** | **Metric** | **Limb** | **Statistical Test** | **Statistic** | **p value** | **Sidak Multiple**  **Comparison** | | **p value** |
| --- | --- | --- | --- | --- | --- | --- | --- | --- | --- |
| Developmental Digigait | ***WT***  N=30  ***AS***  N=20 | Stance Width | Fore | Repeated Measures Mixed effects  Model | Genotype F (1,48) = 22.02  Day F (4.65,222.5) = 22.17  Interaction F (7,335) = 2.076 | **p <.0001***  **p <.0001***  **p =.0456*** | ***WT*** vs  ***AS*** | 22 | p =.7436 |
|  |  |  |  |  |  |  |  | 30 | p =.6105 |
|  |  |  |  |  |  |  |  | 37 | **p =.0002*** |
|  |  |  |  |  |  |  |  | 45 | p =.0562† |
|  |  |  |  |  |  |  |  | 60 | **p =.0018*** |
|  |  |  |  |  |  |  |  | 90 | p =.5782 |
|  |  |  |  |  |  |  |  | 120 | **p =.0178*** |
|  |  |  |  |  |  |  |  | 180 | p =.8457 |
|  |  |  | Hind | Repeated Measures Mixed effects  Model | Genotype F (1,48) = 11.65  Day F (3.197,153) = 220.3  Interaction F (7,335) = 2.912 | **p =.0013***  **p <.0001***  **p =.0057*** |  | 22 | p =.9788 |
|  |  |  |  |  |  |  |  | 30 | p =.1446 |
|  |  |  |  |  |  |  |  | 37 | **p =.0001*** |
|  |  |  |  |  |  |  |  | 45 | p =.5308 |
|  |  |  |  |  |  |  |  | 60 | p =.1303 |
|  |  |  |  |  |  |  |  | 90 | p =.9984 |
|  |  |  |  |  |  |  |  | 120 | p =.8778 |
|  |  |  |  |  |  |  |  | 180 | p =.0792† |
|  |  | Step Length | Fore | Repeated Measures Mixed effects  Model | Genotype F (1,48) = 75.78  Day F (4.772,228.4) = 36.26  Interaction F (7,224) = 6.643 | **p <.0001***  **p <.0001***  **p <.0001*** |  | 22 | p =.3052 |
|  |  |  |  |  |  |  |  | 30 | **p =.0002*** |
|  |  |  |  |  |  |  |  | 37 | **p <.0001*** |
|  |  |  |  |  |  |  |  | 45 | **p <.0001*** |
|  |  |  |  |  |  |  |  | 60 | **p <.0001*** |
|  |  |  |  |  |  |  |  | 90 | **p <.0001*** |
|  |  |  |  |  |  |  |  | 120 | **p <.0001*** |
|  |  |  |  |  |  |  |  | 180 | **p <.0001*** |
|  |  |  | Hind | Repeated Measures Mixed effects  Model | Genotype F (1,48) = 76.48  Day F (4.345, 207.9) = 37.05  Interaction F (7,335) = 6.643 | **p <.0001***  **p <.0001***  **p <.0001*** |  | 22 | **p =.0391*** |
|  |  |  |  |  |  |  |  | 30 | **p =.0003*** |
|  |  |  |  |  |  |  |  | 37 | **p <.0001*** |
|  |  |  |  |  |  |  |  | 45 | **p <.0001*** |
|  |  |  |  |  |  |  |  | 60 | **p <.0001*** |
|  |  |  |  |  |  |  |  | 90 | **p <.0001*** |
|  |  |  |  |  |  |  |  | 120 | **p <.0001*** |
|  |  |  |  |  |  |  |  | 180 | **p =.0002*** |
|  |  | Step Frequency | Fore | Repeated Measures Mixed effects  Model | Genotype F (1,48) = 63.75  Day F (3.113,149) = 25.18  Interaction F (7,335) = 2.834 | **p <.0001***  **p <.0001***  **p =.0070*** |  | 22 | p =.5062 |
|  |  |  |  |  |  |  |  | 30 | **p =.0002*** |
|  |  |  |  |  |  |  |  | 37 | **p <.0001*** |
|  |  |  |  |  |  |  |  | 45 | **p <.0001*** |
|  |  |  |  |  |  |  |  | 60 | **p <.0001*** |
|  |  |  |  |  |  |  |  | 90 | **p <.0001*** |
|  |  |  |  |  |  |  |  | 120 | **p <.0001*** |
|  |  |  |  |  |  |  |  | 180 | **p <.0001*** |
|  |  |  | Hind | Repeated Measures Mixed effects  Model | Genotype F (1,48) = 70.80  Day F (4.171,199.6) = 30.40  Interaction F (7,335) = 3.256 | **p <.0001***  **p <.0001***  **p =.0023*** |  | 22 | p =.0557† |
|  |  |  |  |  |  |  |  | 30 | **p =.0002*** |
|  |  |  |  |  |  |  |  | 37 | **p <.0001*** |
|  |  |  |  |  |  |  |  | 45 | **p <.0001*** |
|  |  |  |  |  |  |  |  | 60 | **p <.0001*** |
|  |  |  |  |  |  |  |  | 90 | **p <.0001*** |
|  |  |  |  |  |  |  |  | 120 | **p <.0001*** |
|  |  |  |  |  |  |  |  | 180 | **p <.0001*** |
|  |  | Stride Duration | Fore | Repeated Measures Mixed effects  Model | Genotype F (1,48) = 75.92  Day F (4.725, 225.5) = 36.13  Interaction F (7,334) = 7.637 | **p <.0001***  **p <.0001***  **p <.0001*** |  | 22 | p =.4274 |
|  |  |  |  |  |  |  |  | 30 | **p =.0002*** |
|  |  |  |  |  |  |  |  | 37 | **p <.0001*** |
|  |  |  |  |  |  |  |  | 45 | **p <.0001*** |
|  |  |  |  |  |  |  |  | 60 | **p <.0001*** |
|  |  |  |  |  |  |  |  | 90 | **p <.0001*** |
|  |  |  |  |  |  |  |  | 120 | **p <.0001*** |
|  |  |  |  |  |  |  |  | 180 | **p <.0001*** |
|  |  |  | Hind | Repeated Measures Mixed effects  Model | Genotype F (1,48) = 76.54  Day F (4.301, 205.2) = 36.38  Interaction F (7,334) = 6.718 | **p <.0001***  **p <.0001***  **p <.0001*** |  | 22 | p =.0573† |
|  |  |  |  |  |  |  |  | 30 | **p =.0002*** |
|  |  |  |  |  |  |  |  | 37 | **p <.0001*** |
|  |  |  |  |  |  |  |  | 45 | **p <.0001*** |
|  |  |  |  |  |  |  |  | 60 | **p <.0001*** |
|  |  |  |  |  |  |  |  | 90 | **p <.0001*** |
|  |  |  |  |  |  |  |  | 120 | **p <.0001*** |
|  |  |  |  |  |  |  |  | 180 | **p =.0002*** |
|  |  | Swing Duration | Fore | Repeated Measures Mixed effects  Model | Genotype F (1,48) = 84.17  Day F (5.103,244.2) = 15.25  Interaction F (7,335) = 3.256 | **p <.0001***  **p <.0001***  **p =.0010*** |  | 22 | p =.2434 |
|  |  |  |  |  |  |  |  | 30 | **p =.0008*** |
|  |  |  |  |  |  |  |  | 37 | **p <.0001*** |
|  |  |  |  |  |  |  |  | 45 | **p <.0001*** |
|  |  |  |  |  |  |  |  | 60 | **p <.0001*** |
|  |  |  |  |  |  |  |  | 90 | **p <.0001*** |
|  |  |  |  |  |  |  |  | 120 | **p <.0001*** |
|  |  |  |  |  |  |  |  | 180 | **p =.0104*** |
|  |  |  | Hind | Repeated Measures Mixed effects  Model | Genotype F (1,48) = 64.94  Day F (5.353,255.4) = 4.387  Interaction F (7,334) = 4.534 | **p <.0001***  **p =.0005***  **p <.0001*** |  | 22 | **p =.0344*** |
|  |  |  |  |  |  |  |  | 30 | **p =.0005*** |
|  |  |  |  |  |  |  |  | 37 | **p <.0001*** |
|  |  |  |  |  |  |  |  | 45 | **p <.0001*** |
|  |  |  |  |  |  |  |  | 60 | **p <.0001*** |
|  |  |  |  |  |  |  |  | 90 | **p =.0002*** |
|  |  |  |  |  |  |  |  | 120 | **p =.0003*** |
|  |  |  |  |  |  |  |  | 180 | **p =.0085*** |
|  |  | Stance Duration | Fore | Repeated Measures Mixed effects  Model | Genotype F (1,48) = 55.22  Day F (4.767,226.8) = 37.3  Interaction F (7,333) = 6.864 | **p <.0001***  **p <.0001***  **p <.0001*** |  | 22 | p =.3019 |
|  |  |  |  |  |  |  |  | 30 | **p =.0015*** |
|  |  |  |  |  |  |  |  | 37 | **p <.0001*** |
|  |  |  |  |  |  |  |  | 45 | **p =.0011*** |
|  |  |  |  |  |  |  |  | 60 | **p <.0001*** |
|  |  |  |  |  |  |  |  | 90 | **p <.0001*** |
|  |  |  |  |  |  |  |  | 120 | **p <.0001*** |
|  |  |  |  |  |  |  |  | 180 | **p <.0001*** |
|  |  |  | Hind | Repeated Measures Mixed effects  Model | Genotype F (1,48) = 55.03  Day F (4.031, 19.2.3) = 72.11  Interaction F (7,334) = 6.857 | **p <.0001***  **p <.0001***  **p <.0001*** |  | 22 | p =.7438 |
|  |  |  |  |  |  |  |  | 30 | **p =.0079*** |
|  |  |  |  |  |  |  |  | 37 | **p <.0001*** |
|  |  |  |  |  |  |  |  | 45 | **p <.0001*** |
|  |  |  |  |  |  |  |  | 60 | **p <.0001*** |
|  |  |  |  |  |  |  |  | 90 | **p <.0001*** |
|  |  |  |  |  |  |  |  | 120 | **p <.0001*** |
|  |  |  |  |  |  |  |  | 180 | **p <.0001*** |
|  |  | Propulsion Duration | Fore | Repeated Measures Mixed effects  Model | Genotype F (1,48) = 34.07  Day F (5.089,242.1) = 69.17  Interaction F (7,333) = 2.297 | **p <.0001***  **p <.0001***  **p =.0268*** |  | 22 | p =.0966† |
|  |  |  |  |  |  |  |  | 30 | **p =.0012*** |
|  |  |  |  |  |  |  |  | 37 | **p =.0213*** |
|  |  |  |  |  |  |  |  | 45 | p =.0684† |
|  |  |  |  |  |  |  |  | 60 | **p =.0335*** |
|  |  |  |  |  |  |  |  | 90 | **p =.0002*** |
|  |  |  |  |  |  |  |  | 120 | **p =.0002*** |
|  |  |  |  |  |  |  |  | 180 | **p =.0012*** |
|  |  |  | Hind | Repeated Measures Mixed effects  Model | Genotype F (1,48) = 58.53  Day F (4.641,221.4) = 64.47  Interaction F (7,334) = 4.26 | **p <.0001***  **p <.0001***  **p =.0002*** |  | 22 | p =.0838† |
|  |  |  |  |  |  |  |  | 30 | **p =.0009*** |
|  |  |  |  |  |  |  |  | 37 | **p <.0001*** |
|  |  |  |  |  |  |  |  | 45 | **p <.0001*** |
|  |  |  |  |  |  |  |  | 60 | **p <.0001*** |
|  |  |  |  |  |  |  |  | 90 | **p <.0001*** |
|  |  |  |  |  |  |  |  | 120 | **p =.0014*** |
|  |  |  |  |  |  |  |  | 180 | **p =.0001*** |
|  |  | Brake Duration | Fore | Repeated Measures Mixed effects  Model | Genotype F (1,48) = 29.31  Day F (5.265, 251.2) = 20.41  Interaction F (7,334) = 4.253 | **p <.0001***  **p <.0001***  **p =.0002*** |  | 22 | p =.9934 |
|  |  |  |  |  |  |  |  | 30 | p >.9999 |
|  |  |  |  |  |  |  |  | 37 | **p =.0038*** |
|  |  |  |  |  |  |  |  | 45 | **p =.0411*** |
|  |  |  |  |  |  |  |  | 60 | **p <.0001*** |
|  |  |  |  |  |  |  |  | 90 | p =.0536† |
|  |  |  |  |  |  |  |  | 120 | p =.0848† |
|  |  |  |  |  |  |  |  | 180 | p =.1315 |
|  |  |  | Hind | Repeated Measures Mixed effects  Model | Genotype F (1,48) = 4.599  Day F (5.455,260.3) = 33.94  Interaction F (7,334) = 1.529 | **p =.0371***  **p <.0001***  p =.15663 |  | 22 | p =.2878 |
|  |  |  |  |  |  |  |  | 30 | p =.7919 |
|  |  |  |  |  |  |  |  | 37 | p =.3263 |
|  |  |  |  |  |  |  |  | 45 | p =.5093 |
|  |  |  |  |  |  |  |  | 60 | p =.5615 |
|  |  |  |  |  |  |  |  | 90 | p >.9999 |
|  |  |  |  |  |  |  |  | 120 | p =.9210 |
|  |  |  |  |  |  |  |  | 180 | p =.8541 |
|  |  | % Shared Stance Time | Hind | Repeated Measures Mixed effects  Model | Genotype F (1,48) = 10.74  Day F (5.919,283.2) = 19.76  Interaction F (7,335) = 3.005 | **p =.0020***  **p <.0001***  **p =.0045*** |  | 22 | p =.6256 |
|  |  |  |  |  |  |  |  | 30 | p >.9999 |
|  |  |  |  |  |  |  |  | 37 | p =.0688† |
|  |  |  |  |  |  |  |  | 45 | **p =.0001*** |
|  |  |  |  |  |  |  |  | 60 | p =.2646 |
|  |  |  |  |  |  |  |  | 90 | p =.0521† |
|  |  |  |  |  |  |  |  | 120 | p =.9109 |
|  |  |  |  |  |  |  |  | 180 | p >.9999 |

| **Test** | **# of Animals** | **Metric** | **Statistical Test** | **Statistic** | **p value** | **Sidak Multiple**  **Comparison** | | **p value** |
| --- | --- | --- | --- | --- | --- | --- | --- | --- |
| Developmental Milestones | ***WT***  N=15  ***AS***  N=19 | Weight | Two Way Repeated Measures ANOVA | Genotype F (1,32) = 0.5838  Day F (2.636,84.34) = 738.5  Interaction F (7,224) = 1.227 | p =.4504  **p <.0001***  p =.2890 | ***WT*** vs  ***AS*** | PND 6 | p =.9617 |
|  |  |  |  |  |  |  | PND 8 | p =.9910 |
|  |  |  |  |  |  |  | PND 10 | p =.9682 |
|  |  |  |  |  |  |  | PND 12 | p =.9610 |
|  |  |  |  |  |  |  | PND 14 | p =.9249 |
|  |  |  |  |  |  |  | PND 16 | p =.9411 |
|  |  |  |  |  |  |  | PND 18 | p =.9782 |
|  |  |  |  |  |  |  | PND 21 | p >.9999 |
|  |  | Body Length | Two Way Repeated Measures ANOVA | Genotype F (1,32) = 1.33  Day F (3.957,126.6) = 375.3  Interaction F (6,192) = 0.5458 | p =.2573  **p <.0001***  p =.7729 |  | PND 6 | p =.9833 |
|  |  |  |  |  |  |  | PND 8 | p =.9401 |
|  |  |  |  |  |  |  | PND 10 | p =.5744 |
|  |  |  |  |  |  |  | PND 12 | p =.6176 |
|  |  |  |  |  |  |  | PND 14 | p >.9999 |
|  |  |  |  |  |  |  | PND 16 | p >.9999 |
|  |  |  |  |  |  |  | PND 18 | p =.9985 |
|  |  | Circle Transverse | Two Way Repeated Measures ANOVA | Genotype F (1,32) = 5.971  Day F (3.894,124.6) = 8.483  Interaction F (6,192) = 1.724 | **p = .0202***  **p <.0001***  p =.1174 |  | PND 6 | p = .9397 |
|  |  |  |  |  |  |  | PND 8 | p =.9421 |
|  |  |  |  |  |  |  | PND 10 | p = .9977 |
|  |  |  |  |  |  |  | PND 12 | p =.8558 |
|  |  |  |  |  |  |  | PND 14 | p >.9999 |
|  |  |  |  |  |  |  | PND 16 | p =.6501 |
|  |  |  |  |  |  |  | PND 18 | p =.1784 |

| **Test** | **# of Animals** | **Metric** | **Statistical Test** | **Statistic** | **p value** | **Sidak Multiple**  **Comparison** | | **p value** |
| --- | --- | --- | --- | --- | --- | --- | --- | --- |
| Replicated-  Independent Cohort  Adult Digigait | ***WT***  N=24  ***AS***  N=17 | Stance Width | Two Way Repeated Measures ANOVA | Genotype F (1,39) = 12.21  Limb F (1,39) = 406.9  Interaction F (1,39) = 1.508 | **p =.0012***  **p <.0001***  p =.2267 | ***WT*** vs  ***AS*** | Forelimb | **p=.0013*** |
|  |  |  |  |  |  |  | Hindlimb | p = .0774† |
|  |  | Step Length | Two Way Repeated Measures ANOVA | Genotype F (1,39) = 25.01  Limb F (1,39) = 4.78  Interaction F (1,39) = 3.003 | **p =.0001***  **p =.0347***  p =.0910† |  | Forelimb | **p<.0001*** |
|  |  |  |  |  |  |  | Hindlimb | **p<.0001*** |
|  |  | Step Frequency | Two Way Repeated Measures ANOVA | Genotype F (1,39) = 26.02  Limb F (1,39) = 2.131  Interaction F (1,39) = 2.592 | **p <.0001***  p =.1523  p =.1155 |  | Forelimb | **p<.0001*** |
|  |  |  |  |  |  |  | Hindlimb | **p<.0001*** |
|  |  | Stride Duration | Two Way Repeated Measures ANOVA | Genotype F (1,39) = 25.15  Limb F (1,39) = 4.297  Interaction F (1,39) = 2.647 | **p <.0001***  **p = .0448***  p =.1118 |  | Forelimb | **p = .0003*** |
|  |  |  |  |  |  |  | Hindlimb | **p = .0003*** |
|  |  | Swing Duration | Two Way Repeated Measures ANOVA | Genotype F (1,39) = 18.26  Limb F (1,39) = 20.49  Interaction F (1,39) = .00326 | **p <.0001***  **p <.0001***  p =.9547 |  | Forelimb | **p = .0285*** |
|  |  |  |  |  |  |  | Hindlimb | **p = .0003*** |
|  |  | Stance Duration | Two Way Repeated Measures ANOVA | Genotype F (1,39) = 14.29  Limb F (1,39) = 24.87  Interaction F (1,39) = 1.972 | **p = .0005***  **p <.0001***  p =.1681 |  | Forelimb | **p<.0001*** |
|  |  |  |  |  |  |  | Hindlimb | **p<.0001*** |
|  |  | Propulsion Duration | Two Way Repeated Measures ANOVA | Genotype F (1,39) = 15.42  Limb F (1,39) = 227.7  Interaction F (1,39) = 0.2848 | **p = .0003***  **p <.0001***  p =.5966 |  | Forelimb | **p = .0174*** |
|  |  |  |  |  |  |  | Hindlimb | **p = .0023*** |
|  |  | Brake Duration | Two Way Repeated Measures ANOVA | Genotype F (1,39) = .03909  Limb F (1,39) = 126.2  Interaction F (1,39) = 0.2335 | p =.8443  **p <.0001***  p =.6317 |  | Forelimb | p = .8524 |
|  |  |  |  |  |  |  | Hindlimb | p = .9572 |

| **Test** | **# of Animals** | **Metric** | **Statistical Test** | **Statistic** | **p value** | **Tukey Multiple**  **Comparison**  **Male vs Female** | | **p value** |
| --- | --- | --- | --- | --- | --- | --- | --- | --- |
| PND60  By Sex  Adult Digigait | ***WT F***  N=13  ***WT M***  N= 17  ***AS F***  N=6  ***AS M***  N=14 | Stance Width | Three Way Repeated Measures ANOVA | Genotype F (1,46) = 14.92  Limb F (1,46) = 676.1  Sex (1,46) = 2.120  Limb x Sex F (1,46) = 0.48  Limb x Genotype F (1,46) = 4.115  Sex x Genotype F (1,46) = 0.1816  Limb x Sex x Genotype F (1,46) = 0.797 | **p <.0001***  **p <.0001***  p =.1522  p =.4919  **p =.0483***  p =.6720  p =.7789 | Forelimb | WT | p=.9939 |
|  |  |  |  |  |  |  | AS | p > .9999 |
|  |  |  |  |  |  | Hindlimb | WT | p=.6633 |
|  |  |  |  |  |  |  | AS | p=.9975 |
|  |  | Step Length | Three Way Repeated Measures ANOVA | Genotype F (1,46) = 43.43  Limb F (1,46) = 5.292  Sex (1,46) = 2.016  Limb x Sex F (1,46) = 2.872  Limb x Genotype F (1,46) = .06910  Sex x Genotype F (1,46) = .008136  Limb x Sex x Genotype F (1,46) = 2.368 | **p <.0001***  **p =.0260***  p =.1624  p =.0969†  p =.7938  p =.9285  p =.1307 | Forelimb | WT | p=.9118 |
|  |  |  |  |  |  |  | AS | p=.9448 |
|  |  |  |  |  |  | Hindlimb | WT | p=.9251 |
|  |  |  |  |  |  |  | AS | p=.9996 |
|  |  | Step Frequency | Three Way Repeated Measures ANOVA | Genotype F (1,46) = 40.09  Limb F (1,46) = 2.493  Sex (1,46) = 2.722  Limb x Sex F (1,46) = 1.737  Limb x Genotype F(1,46) = 0.4039  Sex x Genotype F (1,46) = 0.2173  Limb x Sex x Genotype F (1,46) = 0.05635 | **p <.0001***  p =.1212  **p <.0001***  p =.1940  p =.5282  p =.6433  p =.8134 | Forelimb | WT | p=.5615 |
|  |  |  |  |  |  |  | AS | p=.9847 |
|  |  |  |  |  |  | Hindlimb | WT | p=.7704 |
|  |  |  |  |  |  |  | AS | p=.9993 |
|  |  | Stride Duration | Three Way Repeated Measures ANOVA | Genotype F (1,46) = 43.20  Limb F (1,46) = 4.968  Sex (1,46) = 2.054  Limb x Sex F (1,46) = 2.768  Limb x Genotype F(1,46) = 0.03376  Sex x Genotype F (1,46) = 0.00739  Limb x Sex x Genotype F (1,46) = 2.19 | **p <.0001***  **p =.0308***  p =.1586  p =.1029  p =.8550  p =.9318  p =.1458 | Forelimb | WT | p=.9079 |
|  |  |  |  |  |  |  | AS | p=.9449 |
|  |  |  |  |  |  | Hindlimb | WT | p=.9238 |
|  |  |  |  |  |  |  | AS | p=.9995 |
|  |  | Swing Duration | Three Way Repeated Measures ANOVA | Genotype F (1,46) = 37.60  Limb F (1,46) = 69.11  Sex (1,46) = 0.0007  Limb x Sex F (1,46) = 10.77  Limb x Genotype F(1,46) = 1.484  Sex x Genotype F (1,46) = 0.1231  Limb x Sex x Genotype F (1,46) = 0.0017 | **p <.0001***  **p <.0001***  p =.9789  **p =.0020***  p =.2294  p =.7273  p =.9672 | Forelimb | WT | p > .9999 |
|  |  |  |  |  |  |  | AS | p=.9973 |
|  |  |  |  |  |  | Hindlimb | WT | p=.9785 |
|  |  |  |  |  |  |  | AS | p > .9999 |
|  |  | Stance Duration | Three Way Repeated Measures ANOVA | Genotype F (1,46) = 38.56  Limb F (1,46) = 61.25  Sex (1,46) = 6.493  Limb x Sex F (1,46) = 0.5569  Limb x Genotype F(1,46) = 1.123  Sex x Genotype F (1,46) = 0.1872  Limb x Sex x Genotype F (1,46) = 2.032 | **p <.0001***  **p <.0001***  **p =.0142***  p =.4593  p =.2948  p =.6673  p =.1608 | Forelimb | WT | p=.5985 |
|  |  |  |  |  |  |  | AS | p=.8587 |
|  |  |  |  |  |  | Hindlimb | WT | p=.0858† |
|  |  |  |  |  |  |  | AS | p=.9456 |
|  |  | Propulsion  Duration | Three Way Repeated Measures ANOVA | Genotype F (1,46) = 26.48  Limb F (1,46) = 560.9  Sex (1,46) = 7.836  Limb x Sex F (1,46) = 2.411  Limb x Genotype F(1,46) = 24.83  Sex x Genotype F (1,46) = 0.09822  Limb x Sex x Genotype F (1,46) = 0.2994 | **p <.0001***  **p <.0001***  **p =.0075***  p =.1273  **p <.0001***  p =.7554  p =.5869 | Forelimb | WT | p=.7541 |
|  |  |  |  |  |  |  | AS | p=.9462 |
|  |  |  |  |  |  | Hindlimb | WT | p=.0531† |
|  |  |  |  |  |  |  | AS | p=.7016 |
|  |  | Brake Duration | Three Way Repeated Measures ANOVA | Genotype F (1,46) = 7.394  Limb F (1,46) = 238.3  Sex (1,46) = 0.0006998  Limb x Sex F (1,46) = 0.8052  Limb x Genotype F(1,46) = 24.00  Sex x Genotype F (1,46) = 0.07782  Limb x Sex x Genotype F (1,46) = 0.1849 | **p =.0092***  **p <.0001***  p =.9790  p =.3742  **p <.0001***  p =.7815  p =.6692 | Forelimb | WT | p=.9998 |
|  |  |  |  |  |  |  | AS | p > .9999 |
|  |  |  |  |  |  | Hindlimb | WT | p > .9999 |
|  |  |  |  |  |  |  | AS | p=.9979 |

| **Test** | **# of Animals** | **Metric** | **Statistical Test** | **Statistic** | **p value** | **Sidak Multiple**  **Comparison** | | **p value** |
| --- | --- | --- | --- | --- | --- | --- | --- | --- |
| Adult Digigait  18 cm/s | ***WT***  N=16  ***AS***  N=15 | Stance Width | Two Way Repeated Measures ANOVA | Genotype F (1,29) = 9.114  Limb F (1,29) = 529.9  Interaction F (1,29) = 0.9944 | **p =.0052***  **p <.0001***  p =.3269 | ***WT*** vs  ***AS*** | Forelimb | **p=.0077*** |
|  |  |  |  |  |  |  | Hindlimb | p = .1462 |
|  |  | Step Length | Two Way Repeated Measures ANOVA | Genotype F (1,29) = 11.52  Limb F (1,29) = 7.396  Interaction F (1,29) = 0.01483 | **p =.0020***  **p =.0109***  p =.9039 |  | Forelimb | **p=.0033*** |
|  |  |  |  |  |  |  | Hindlimb | **p=.0040*** |
|  |  | Step Frequency | Two Way Repeated Measures ANOVA | Genotype F (1,29) = 12.18  Limb F (1,29) = 5.359  Interaction F (1,29) = 0.5354 | **p =.0016***  **p =.0279***  p =.4702 |  | Forelimb | **p=.0016*** |
|  |  |  |  |  |  |  | Hindlimb | **p=.0065*** |
|  |  | Stride Duration | Two Way Repeated Measures ANOVA | Genotype F (1,29) = 11.45  Limb F (1,29) = 7.537  Interaction F (1,29) = 0.0719 | **p = .0021***  **p = .0103***  p =.7904 |  | Forelimb | **p = .0030*** |
|  |  |  |  |  |  |  | Hindlimb | **p = .0045*** |
|  |  | Swing Duration | Two Way Repeated Measures ANOVA | Genotype F (1,29) = 8.184  Limb F (1,29) = 14.52  Interaction F (1,29) = 0.0259 | **p = .0078***  **p = .0007***  p =.8730 |  | Forelimb | **p = .0288*** |
|  |  |  |  |  |  |  | Hindlimb | **p = .0202*** |
|  |  | Stance Duration | Two Way Repeated Measures ANOVA | Genotype F (1,29) = 10.97  Limb F (1,29) = 31.31  Interaction F (1,29) = 0.1235 | **p = .0025***  **p <.0001***  p =.7872 |  | Forelimb | **p = .0050*** |
|  |  |  |  |  |  |  | Hindlimb | **p = .0113*** |
|  |  | Propulsion Duration | Two Way Repeated Measures ANOVA | Genotype F (1,29) = 10.71  Limb F (1,29) = 206.8  Interaction F (1,29) = 3.443 | **p = .0028***  **p <.0001***  p =.0737† |  | Forelimb | p = .1867 |
|  |  |  |  |  |  |  | Hindlimb | **p = .0008*** |
|  |  | Brake Duration | Two Way Repeated Measures ANOVA | Genotype F (1,29) = 2.027  Limb F (1,29) = 104.9  Interaction F (1,29) = 5.960 | p =.1652  **p <.0001***  **p = .0210*** |  | Forelimb | **p = .0235*** |
|  |  |  |  |  |  |  | Hindlimb | p = .9389 |

| **Test** | **# of Animals** | **Metric** | **Statistical Test** | **Statistic** | **p value** | **Sidak Multiple**  **Comparison** | | **p value** |
| --- | --- | --- | --- | --- | --- | --- | --- | --- |
| Adult Digigait  36 cm/s | ***WT***  N=16  ***AS***  N=15 | Stance Width | Two Way Repeated Measures ANOVA | Genotype F (1,30) = 5.146  Limb F (1,30) = 265.1  Interaction F (1,30) = 0.1019 | **p =.0307***  **p <.0001***  p =.7517 | ***WT*** vs  ***AS*** | Forelimb | p = .3235 |
|  |  |  |  |  |  |  | Hindlimb | p = .1422 |
|  |  | Step Length | Two Way Repeated Measures ANOVA | Genotype F (1,30) = 3.794  Limb F (1,30) = 0.3506  Interaction F (1,30) = 0.6458 | p =.0609†  p =.5582  p =.4280 |  | Forelimb | p = .3111 |
|  |  |  |  |  |  |  | Hindlimb | p = .0779† |
|  |  | Step Frequency | Two Way Repeated Measures ANOVA | Genotype F (1,30) = 5.366  Limb F (1,30) = 0.3027  Interaction F (1,30) = 0.3698 | **p =.0275***  p =.5862  p =.5477 |  | Forelimb | p = .1566 |
|  |  |  |  |  |  |  | Hindlimb | **p = .0453*** |
|  |  | Stride Duration | Two Way Repeated Measures ANOVA | Genotype F (1,30) = 3.877  Limb F (1,30) = 0.3780  Interaction F (1,30) = 0.8506 | p = .0582†  p =.5433  p =.3638 |  | Forelimb | p = .3252 |
|  |  |  |  |  |  |  | Hindlimb | p = .0662† |
|  |  | Swing Duration | Two Way Repeated Measures ANOVA | Genotype F (1,30) = 1.189  Limb F (1,30) = 4.114  Interaction F (1,30) = 0.8081 | p = .2842  p =.0515†  p =.3758 |  | Forelimb | p = .9490 |
|  |  |  |  |  |  |  | Hindlimb | p = .2991 |
|  |  | Stance Duration | Two Way Repeated Measures ANOVA | Genotype F (1,30) = 3.396  Limb F (1,30) = 4.881  Interaction F (1,30) = 0.000 | p =.0753†  **p = .0349***  p > .9999 |  | Forelimb | p = .3030 |
|  |  |  |  |  |  |  | Hindlimb | p = .3030 |
|  |  | Propulsion Duration | Two Way Repeated Measures ANOVA | Genotype F (1,30) = 3.139  Limb F (1,30) = 105.0  Interaction F (1,30) = 0.6079 | p =.0866†  **p <.0001***  p =.4417 |  | Forelimb | p = .6380 |
|  |  |  |  |  |  |  | Hindlimb | p = .1319 |
|  |  | Brake Duration | Two Way Repeated Measures ANOVA | Genotype F (1,30) = 0.07868  Limb F (1,30) = 90.69  Interaction F (1,30) = 0.8755 | p =.7810  **p <.0001***  p =.3569 |  | Forelimb | p = .7061 |
|  |  |  |  |  |  |  | Hindlimb | p = .9525 |
